# Strong positive selection in *Aedes aegypti* and the rapid evolution of insecticide resistance

**DOI:** 10.1101/2022.07.27.501751

**Authors:** R. Rebecca Love, Josh R. Sikder, Rafael J. Vivero, Daniel R. Matute, Daniel R. Schrider

## Abstract

*Aedes aegypti* vectors the pathogens that cause dengue, yellow fever, Zika virus, and chikungunya, and is a serious threat to public health in tropical regions. Decades of work has illuminated many aspects of *Ae. aegypti*’s biology and global population structure, and has identified insecticide resistance genes; however, the size and repetitive nature of the *Ae. aegypti* genome have limited our ability to detect positive selection in this mosquito. Combining new whole genome sequences from Colombia with publicly available data from Africa and the Americas, we identify multiple strong candidate selective sweeps in *Ae. aegypti*, many of which overlap genes linked to or implicated in insecticide resistance. We examine the voltage-gated sodium channel gene in three American cohorts, and find evidence for successive selective sweeps in Colombia. The most recent sweep encompasses an intermediate-frequency haplotype containing four candidate insecticide resistance mutations that are in near-perfect linkage disequilibrium with one another in the Colombian sample. We hypothesize that this haplotype may continue to rapidly increase in frequency and perhaps spread geographically in the coming years. These results extend our knowledge of how insecticide resistance has evolved in this species, and add to a growing body of evidence suggesting *Ae. aegypti* has an extensive genomic capacity to rapidly adapt to insecticide-based vector control.

## INTRODUCTION

The mosquito *Aedes (Stegomyia) aegypti* (Linnaeus, 1762) vectors the arboviruses causing dengue, Zika, yellow fever, and chikungunya (Viglietta et al. 2021), the former two of which have already caused pandemics in the first two decades of the 21^st^ century. No vaccine is publicly available for chikungunya or Zika, and treatment for all four diseases is generally supportive or symptomatic only. While considerable uncertainty exists about the actual number of annual dengue cases worldwide (Wilder-Smith and Byass 2016, Bhatt et al. 2013), one common estimate suggests approximately 400 million cases a year as of 2010 (Bhatt et al. 2013).

Because of its importance to public health, as well as its facility as a model organism, *Ae. aegypti* has been extensively studied. Two subspecies or forms have been identified, initially distinguished on the basis of morphology with corresponding differences in behavior (Mattingly 1957), though the morphological characters in question have long been recognized as subject to variation (Mattingly 1957, Brown et al. 2011). *Ae. aegypti formosus*, recognizable by its dark abdomen, generally prefers forest and other outdoor habitats, and is limited to Africa; in contrast, *Ae. aegypti aegypti* has pale scales on the first abdominal tergite, is found widely across tropical regions of the world outside of Africa, is highly anthropophilic, and is found in close association in humans, often resting indoors and ovipositing in anthropogenic habitats (Mattingly 1957). The two subspecies are genetically distinct as well (Tabachnick and Powell 1979, Gloria-Soria, Ayala, et al. 2016, Crawford et al. 2017, Kotsakiozi, Evans, et al. 2018), though both morphological and genetic distinctions break down in some parts of West Africa (Gloria-Soria, Ayala, et al. 2016, Crawford et al. 2017, Kotsakiozi, Evans, et al. 2018). In addition, the two forms may be differentially susceptible to dengue (Sylla et al. 2009), Zika virus (Aubry et al. 2020), and yellow fever (Tabachnick et al. 1985), though the latter phenotype appears to be highly variable depending on mosquito and viral strain (Tabachnick et al. 1985, Dickson et al. 2014). A third form, *queenslandensis*, was initially recognized on the basis of morphology (Mattingly 1957) but appears to represent morphological variation of *Ae. aegypti aegypti* (Rašić et al. 2016).

*Ae. aegypti* tends to have low unaided dispersal (Harrington et al. 2005), though it may fly further if suitable oviposition sites or opportunities to blood-feed are lacking (Reiter et al. 1995, Codeço et al. 2007). The widespread tropical distribution of *Ae. aegypti aegypti* is therefore attributed to its close association with humans (Christophers 1960). Originating in Africa (Tabachnick 1991, Brown et al. 2011, Brown et al. 2014), *Ae. aegypti* spread to the Americas (Brown et al. 2014, Gloria-Soria, Ayala, et al. 2016), likely due to mass transatlantic trafficking of enslaved people. It arrived in Asia more recently (Tabachnick 1991, Brown et al. 2014), possibly facilitated by the opening of the Suez Canal (Tabachnick 1991, Powell et al. 2018). The non-African domestic populations appear to result from one colonization event (Crawford et al. 2017, Rose et al. 2020), descending from a domesticated subpopulation in western Africa (Crawford et al. 2017) or possibly central Africa (Kotsakiozi, Evans, et al. 2018); human preference is linked to expression and sequence variation in the *Or4* gene (McBride et al. 2014, Rose et al. 2020) as well as dry season intensity (Rose et al. 2020).

Population structure tends to be less pronounced within Africa than outside Africa (Brown et al. 2011, Brown et al. 2014, Gloria-Soria, Ayala, et al. 2016); outside Africa, specimens show strong geographic clustering (Rašić et al. 2014, Evans et al. 2015, Schmidt et al. 2020) but extensive connectivity is seen between some geographically distant populations (Gonçalves da Silva et al. 2012, Monteiro et al. 2014, Gloria-Soria, Ayala, et al. 2016). Populations are often dynamic, showing sometimes extensive changes in genetic composition within a few years (Gloria-Soria, Kellner, et al. 2016), and reinvading areas where they had been eradicated (Kelly et al. 2021) or out-competed (Brennan et al. 2021).

Overall, the global genetic structure of *Ae. aegypti* has been well-studied; however, because of the size and repetitive nature of its genome, nearly all previous work has used either sequences from a small number of genes (Bennett et al. 2016), microsatellites (Brown et al. 2011, Gloria-Soria, Ayala, et al. 2016), mtDNA (Gonçalves da Silva et al. 2012, Bennett et al. 2016), genotyping arrays (Evans et al. 2015, Kotsakiozi, Evans, et al. 2018), or reduced representation sequencing (Brown et al. 2014, Rašić et al. 2014). Exome-based (Crawford et al. 2017, Dickson et al. 2017) and whole genome-based population resequencing have only been performed recently (Lee et al. 2019, Rose et al. 2020, Kelly et al. 2021), and the improvement of the *Ae. aegypti* reference genome to a chromosomal-level assembly (Matthews et al. 2018) will greatly facilitate such efforts moving forward. Studies using a small number of loci are extremely limited in their ability to detect recent positive selection, which skews patterns of diversity across larger genomic regions (Maynard Smith and Haigh 1974, Kaplan et al. 1989, Kim and Nielsen 2004). As a consequence, we know relatively little about the extent of recent positive selection or the broad-scale genomic underpinnings of adaptation in *Ae. aegypti*, compared to our more extensive knowledge of the species’ spread and structure. Positive selection has been observed on genomic regions linked to human preference (Rose et al. 2020), and on genes that enable females to retain eggs during periods of drought (Venkataraman et al. 2022); however, with the exception of a putative selective sweep on an insecticide resistance gene (the *Vgsc* gene, discussed below) identified in a sample from southern Mexico (Saavedra-Rodriguez et al. 2019), we know little about how its genome has responded to these selective pressures at a scale beyond that of individual loci.

In some parts of the Americas, *Ae. aegypti* has been subject to insecticide-based vector control for nearly a century. In the early part of the 20^th^ century, some of the techniques used to target *Ae. aegypti* in Latin America included fumigation of dwellings and “oiling” of water sources to kill larvae (Severo 1955). DDT was used beginning in 1945 (Severo 1955) and extensively thereafter, enabling the campaign for continent-wide eradication in the Americas which began officially in 1947 (Soper 1965). The campaign was initially successful in eradicating *Ae. aegypti* from most of South and Central America (Soper 1965); however, DDT resistance had appeared in the Americas by 1964 (Soper 1965), and for this and other reasons, the campaign faltered. *Ae. aegypti* soon spread across South and Central America again (Jansen and Beebe 2010).

Resistance to DDT in *Ae. aegypti* is conferred by both increased metabolism as well as non-metabolic factors (Rathor and Wood 1981, Hemingway et al. 1989, Brengues et al. 2003). The best-characterized non-metabolic factor is *k*nock*d*own *r*esistance (*kdr*), first identified in houseflies, and conferring resistance to pyrethroids as well (Farnham 1977); *kdr*, caused by point mutations in the voltage-gated sodium channel gene (*Vgsc*, (Williamson et al. 1996)) has also been found in a wide variety of other insects, including *Drosophila* (Pittendrigh et al. 1997), anopheline mosquitoes (Martinez-Torres et al. 1998, Clarkson et al. 2021), tobacco budworms (Taylor et al. 1993), cockroaches (Dong and Scott 1994), and many others (Davies et al. 2007).

Unsurprisingly considering the breadth of distribution of these mutations in other insects, an extensive array of nonsynonymous point mutations associated with pyrethroid resistance have been identified in *Ae. aegypti*. (Because this resistance factor was originally identified in *Musca domestica*, the *Musca domestica* amino acid numberings are often used; to avoid ambiguity, we will follow this convention.) Some, such as F1534C (Kawada et al. 2009, Harris et al. 2010), are globally distributed. Others appear limited to specific continents or regions; V1016I has so far only been recorded from the Americas (Fan et al. 2020) and Africa (Sombié et al. 2019), while V1016G has only been recorded from Asia (Fan et al. 2020). Similarly, S989P appears limited to Asia (Fan et al. 2020), while S723T has only been identified in the Americas (Fan et al. 2020), and V410L has been found in the Americas (Fan et al. 2020) and Africa (Ayres et al. 2020). A subset of the mutations associated with pyrethroid resistance, including F1534C (Hu et al. 2011, Du et al. 2013, Hirata et al. 2014, Haddi et al. 2017), V1016G (Du et al. 2013, Hirata et al. 2014), I1011M (Du et al. 2013) and V410L (Haddi et al. 2017) have been shown experimentally to directly cause pyrethroid resistance.

These mutations are often found in combination with each other; for example, V410L has been observed in conjunction with F1534C (Haddi et al. 2017, Saavedra-Rodriguez et al. 2018) and V1016I (Saavedra-Rodriguez et al. 2018), and S989P is very frequently found on a V1016G background (Kawada et al. 2014, Al Nazawi et al. 2017). Recurrent evolution (Fan et al. 2020) and recombination (Saavedra-Rodriguez et al. 2018) have been suggested as contributing factors to the combinations and distribution of these various mutations. A similar situation of recurrent mutation at the same site, and the presence of haplotypes comprising multiple resistance alleles, has been observed in *An. gambiae* and *An. coluzzii* (Clarkson et al. 2021).

Natural selection associated with insecticide resistance has been widely observed in other insects, including other disease vectors such as *Anopheles gambiae* and *An. coluzzii* (Miles et al. 2017, Xue et al. 2021) and *Anopheles funestus* (Weedall et al. 2020); agricultural pests such as *Leptinotarsa decemlineata* (Pélissié et al. 2022) and *Helicoverpa armigera* (Walsh et al. 2018); and the model organism *Drosophila melanogaster* (Turner et al. 2008, Kolaczkowski et al. 2011), Garud et al. 2015, Schrider et al. 2016). Thus, it seems probable that positive selection for insecticide resistance would be widespread in *Aedes* as well. In addition to selection at specific insecticide resistance loci, vector control programs may cause population contractions (Urdaneta-Marquez and Failloux 2011), which can affect genome-wide change in patterns of diversity. In Colombia, vector control relies on pyrethroids and organophosphates (temephos and malathion) (Aguirre-Obando et al. 2015).

Here, we examine patterns of genetic variation in a set of 30 *Aedes aegypti* genomes collected from Colombia. We also combine these data with previously published *Aedes* genomic data to examine global patterns of genetic variation. Finally, we turn our attention toward detecting signatures of recent positive selection in *Aedes* samples from Africa and the Americas. In five of the six countries examined, we find strong signals of selection in regions containing genes that have been found to be associated with insecticide resistance in *Aedes* or other insects. We discuss the implications of these findings for adaptation to ongoing and future mosquito control efforts.

## RESULTS

### Quality control, alignment, and variant genotyping

We used a stringent filtering approach to control the quality of the sequenced samples (Methods). We removed six specimens from the Colombian sample that had mean coverage below 15x or fewer than 70% of reads mapping to the reference genome, leaving a sample of 10 specimens from Cali and 24 from Río Claro (see Table S1 for complete specimen list).

In addition, we examined publicly available sequence data from the United States of America (*n*=27) from Lee et al. 2019 (Lee et al. 2019), and from Santarém, Brazil (*n*=18), Franceville, Gabon (*n*=13), Kaya Bomu, Kenya (*n*=19), and Ngoye, Senegal (*n*=20) from Rose et al. (2020) (Rose et al. 2020), for a total of six cohorts. The Ngoye cohort comes from a population with a strong preference for human over non-human odor as well as a large “domestic” ancestry component; the Franceville cohort originates from a population with a strong preference for non-human odor, and no domestic ancestry; the Kaya Bomu population is intermediate in both traits (Rose et al. 2020).

After removing the six specimens in our focal Colombian sample that failed our quality filters, mean read depth in each population ranged from 10.16 (USA) to 21.10 (Colombia), with between 94.80% (Gabon) and 98.19% (USA) of reads mapping on average (Table S2).

Variant calling produced an initial data set of 247,108,978 variants in 131 specimens. After filtering, we retained 43,213,550 high-quality single nucleotide polymorphisms (SNPs) with a known location on one of the three autosomes, and an additional 226,687 SNPs in regions of the genome that could not be confidently assigned to chromosomes. Our analysis focused on the SNPs on the three autosomes. The number of SNPs found in each cohort, before and after applying filters based on relatedness as discussed below, is shown in Table S3.

In keeping with previous work, substantially more SNPs were called per specimen in the African sample than in the American sample (Table S3). Within the African sample, the Senegal sample had markedly fewer SNPs called per specimen than the Gabon or Kenya samples, perhaps reflecting that this is a human specialist population (Rose et al. 2020) and therefore may have experienced a bottleneck during the transition towards close proximity with humans (Crawford et al. 2017).

We also filtered genomes based on kinship coefficient after variant calling. From the American sample, ten pairs of specimens, involving 19 specimens (*i.e.*, one specimen appeared in two pairs) had a kinship coefficient greater than 0.2 (see Figure S1 for kinship coefficients between all pairs of specimens). From the African sample, four pairs involving eight specimens had a kinship coefficient greater than 0.05. We removed 13 specimens (four from Colombia, three from Brazil, two from the USA, three from Kenya, and one from Senegal), leaving a total of 118 remaining specimens.

With one exception, all pairs of specimens with a kinship coefficient greater than 0.05 were from the same country. However, one male specimen from Colombia had a kinship coefficient of 0.07 with a Brazilian specimen. While KING-robust may produce biased kinship coefficients in certain cases of admixture (Conomos et al. 2016), this finding is in keeping with previous observations of shared ancestry between South American countries (Monteiro et al. 2014, Kotsakiozi, Gloria-Soria, et al. 2018).

### Patterns of genetic diversity within and between populations

To investigate the extent and patterns of population structure of our data set, we used principal component analysis. The first principal component for each chromosome explained 12-13% of the variance, and generally separated samples from Africa vs. samples from North and South America (Figure 1, Figure S2). The second principal component, explaining 4-5% of the variance, separated samples from North and South America along a cline stretching from the USA sample to the Colombian sample. The two Colombian cohorts were clearly separated, with the Brazilian sample falling in between.

**Figure 1.**
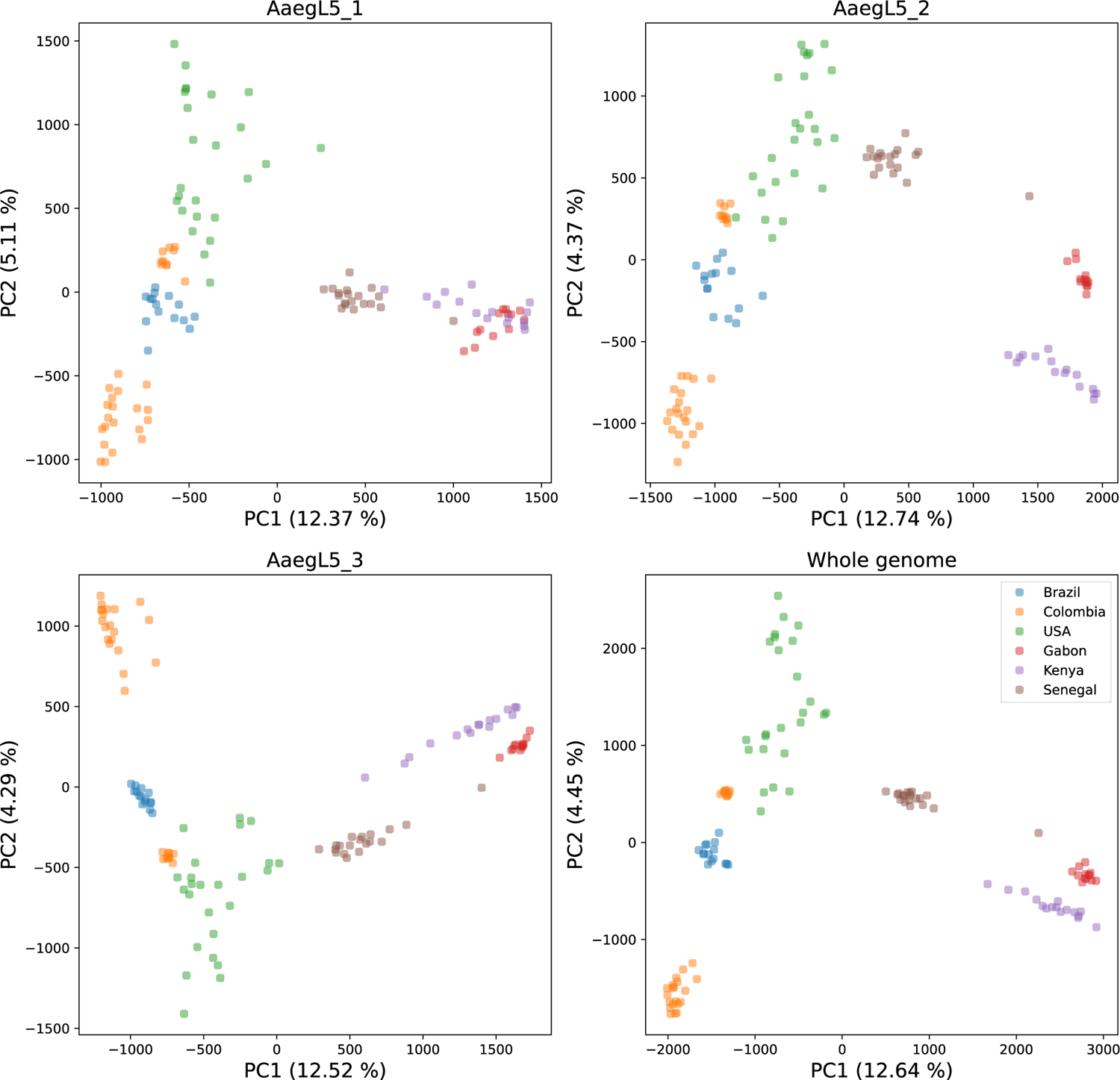
Principal component analysis of all filtered SNPs, showing each chromosome separately (left panels and top-right panel) and combining all three chromosomes (bottom-right panel).

On each chromosome, the Senegal sample was closer in the first principal component to the North and South American samples than were the Gabon and Kenya samples. This is in keeping with previous findings (Crawford et al. 2017, Rose et al. 2020) that have suggested a Senegalese origin of the domestic populations. On chromosome 1, the Kenya and Gabon samples are generally mixed, while they exhibit greater separation on chromosomes 2 and 3. Overall, the separation between the American and African samples is most pronounced on chromosome 1, largely due to the shifting position of the Senegal cohort on the other chromosomes: on chromosome 2, Senegal clusters particularly closely to the USA cohort, and on chromosome 3 Senegal is placed in an intermediate position relative to its position on the other two chromosomes.

Next, we quantified the extent of genetic diversity and the prevalence of rare versus common polymorphisms by measuring nucleotide diversity (π) and Tajima’s *D*. Nucleotide diversity was lower in the American samples than in the African samples (0.0021 vs 0.0031 on average; Figure S3, Table S4), consistent with previous reports of greater allelic richness in African samples (Gloria-Soria, Ayala, et al. 2016). Tajima’s *D* was higher in the American cohorts, consistent with the loss of rare variants caused by a population contraction. In all cohorts, nucleotide diversity tended to be lower near the telomeres, along with reductions in nucleotide diversity and Tajima’s *D* observed near the centromeres (Figure S4). Tajima’s *D* in the African cohorts tended to be lower towards the telomeres, a trend not observed in the American cohorts.

Next, we quantified between-population diversity using *F_ST_*. As expected, mean genome-wide *F_ST_* values were lowest within continents and elevated between continents (Table S5). The decreased *F_ST_* between all American populations and Senegal, compared to American populations and Gabon or Kenya, are consistent with the West African origin of domestic *Ae. aegypti*.

We also measured linkage disequilibrium (LD) within 500 kb windows and found that LD in the Brazilian population was noticeably elevated on all three chromosomes, while the other five populations overlapped more closely (Figure S5, Table S6). Indeed, mean linkage disequilibrium is much higher in the Brazilian sample than in the other five populations—nearly twice as high in Colombia and Gabon, the two populations with the next-highest mean value of *r*^2^ (Table S6).

### Strong signatures of selective sweeps in insecticide resistance loci

We scanned the genome for signatures of recent selective sweeps using SweepFinder2 (DeGiorgio et al. 2016) (Methods). Strong sweep signals were observed in all populations, with the most prominent peaks in SweepFinder2’s composite likelihood ratio (CLR) seen in the samples from Brazil, Colombia, and Gabon (Figure 2). CLR peaks were also observed at the beginnings or ends of chromosomes in all populations except Senegal; the size and shape of these may be a result of the paucity of recombination and the lack of polymorphisms in these regions, respectively.

**Figure 2.**
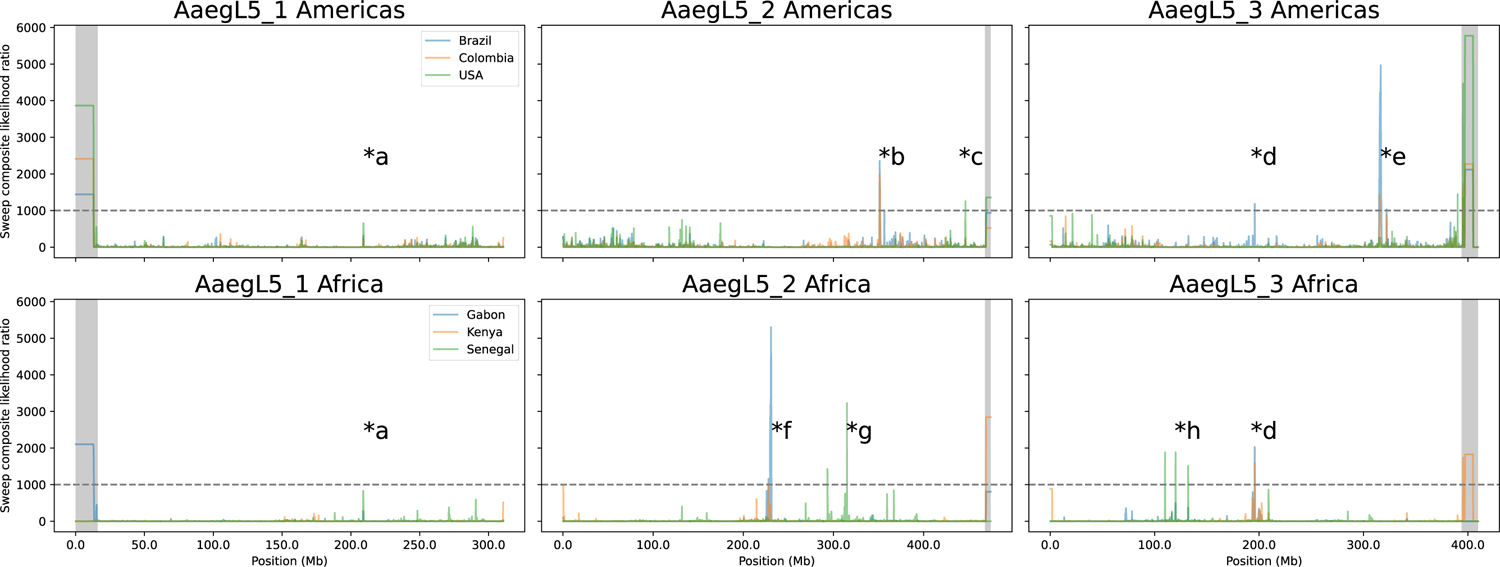
Genome-wide scan of candidate selective sweeps. Shaded regions at the ends of chromosomes indicate putative technical artifacts excluded from identification of outliers. Each * with a letter marks a sweep examined in more detail in the text and in the corresponding panel of Figure S6. Horizontal dashed lines indicate our threshold of CLR=1000, used in identifying sweeps examined further.

We individually examined eight peaks or groups of peaks (marked with stars in Figure 2), that either contained a CLR value greater than 1,000, or had values near 1,000 in both the Americas and Africa. An example of the latter is found on chromosome 1 at approximately 209 Mb (the peak marked “*a” in Figure 2), where all cohorts except Kenya show some signal of a sweep (Fig S6a). This putative swept region overlaps 21 genes (Table S7); 19 of these code for a protein, and six are annotated with a specific predicted function, with experimental evidence available for three. The sulphate transporter gene AAEL012037 is regulated by the hormone 20-hydroxyecdysone via the ecdysone receptor (Roy et al. 2022). Another sulphate transporter, AAEL012041, is upregulated after mating (Camargo et al. 2020). The zinc finger gene AAEL012039, part of the Hedgehog signaling pathway, is upregulated in the brains of female pupae compared to those of male pupae, and localized to the optic lobe (Tomchaney et al. 2014).

Within the American samples, the two most prominent peaks, one each on chromosomes 2 and 3, are shared to some extent by both the Brazilian and the Colombian sample. A peak on chromosome 2 (“*b”), from approximately 351 Mb to 352 Mb, overlaps 27 protein-coding genes and four non-coding RNAs (Figure 2, Fig S6b, Table S7); 18 protein-coding genes have an unspecified functional product, while the genes with specific predicted functions include an ankyrin gene (AAEL013466; discussed below) as well as a cluster of 8 glutathione transferases. Glutathione transferases have a well-known function insecticide resistance, and have been implicated in resistance to several classes of insecticides (Enayati et al. 2005).

A secondary peak is found nearby in the Brazilian sample, but not the Colombian sample, from approximately 356.3 Mb to 356.5 Mb on chromosome 2. There are one tRNA and six protein-coding genes in the vicinity of this peak (Fig S6b, Table S7), three of which have a specific functional prediction. AAEL002055, a neuroendocrine protein, was previously identified as carrying a single nucleotide showing significant spatial structure within a Brazilian population (Rašić et al. 2015); *CYP4H31*/AAEL002085 is a cytochrome P450 identified as overtranscribed in larvae selected to be resistant to the neonicotinoid insecticide imidacloprid (Riaz et al. 2013). Its expression levels are altered in larvae exposed to DDT, malathion and temephos (El-Garj et al. 2016), and it has been implicated in deltamethrin resistance in *Culex* (Li et al. 2021).

Also on chromosome 2, a candidate sweep found only in the USA cohort extends from approximately 446 to 446.5 Mb, (“*c”, Figure 2, Figure S6c). This region overlaps 27 genes, 18 of which are protein-coding and 8 of which have specific functional predictions (Table S7). Of these 18, three are known in the literature. The glutathione transferase *GSTX2* (AAEL010500) is upregulated in response to a variety of exposures, including to the insecticide propoxur (David et al. 2010) and to brackish water (Durant et al. 2021) in larvae, as well as to heme in a cell line and in midguts of females after a blood meal (Bottino-Rojas et al. 2015); *GSTX2* is also constitutively upregulated in a strain resistant to organophosphate and pyrethroid insecticides compared to a susceptible strain (Bottino-Rojas et al. 2018). The protein product of transcriptional regulator *ATRX* (AAEL010502) is found in eggshell preparations (Marinotti et al. 2014), and the fk506-binding protein *FKBP12* (AAEL010491) is upregulated in a population with high rates of *Wolbachia* wMEL infection (Martins et al. 2021).

On chromosome 3, Gabon, Kenya, and Brazil share a sweep at approximately 200 Mb (“*d”, Figure 2). Closer examination reveals a shared peak at 196 Mb, and a peak in Kenya between 202 and 202.5 Mb, the latter of which occurs at much lower magnitude in Gabon (Figure S6d). The former peak, at 196 Mb, overlaps 11 genes, 10 of which are protein-coding and four of which have a specific predicted function. One of these, AAEL007292, is a larval lethal gene involved in the proper development of neural synapses (Hapairai et al. 2017). The latter peak overlaps one protein-coding gene and one noncoding RNA (Table S7).

The highest peak identified in any American sample occurs in the Brazilian sample, near 315 Mb on chromosome 3; this peak also occurs in the Colombian sample at a lesser magnitude (“*e”, Figure 2). Closer investigation shows this peak extending from approximately 314.5 Mb to 318 Mb in the Brazilian sample (Fig S6e). This region overlaps 36 protein-coding genes and 11 non-coding RNAs (Table S7). Of the protein-coding genes, 13 have a specific functional prediction present in VectorBase; we highlight five genes that are known from the literature. The Vanin-like protein AAEL006034 has been identified as potentially involved in metabolism of the insecticide indoxacarb (Smith et al. 2018). The calmodulin gene AAEL012326 is overexpressed in a strain susceptible to dengue compared to one refractory to dengue (Chauhan et al. 2012), and produces a protein product that is found in extracellular vesicles produced by *Ae. aegypti* cells infected by dengue, but not uninfected control cells (Gold et al. 2020). The putative 60S ribosomal protein gene AAEL011587 is upregulated in response to chikungunya infection and to chikungunya and dengue coinfection (Shrinet et al. 2017). The heme peroxidase gene *HPX1* (AAEL006014) is overexpressed in response to pyrethroid (permethrin) selection (Saavedra-Rodriguez et al. 2012).

Finally, this peak near 315 Mb on chromosome 3 also includes *Vgsc* (AAEL023266), which has been extensively associated with insecticide resistance via non-synonymous single nucleotide changes that cause resistance to pyrethroids, DDT, or both (Hu et al. 2011, Du et al. 2013, Hirata et al. 2014, Haddi et al. 2017). In the Brazilian sample, *Vgsc* is roughly centered in the putative sweep, while in our Colombian sample, the region of highest sweep likelihood is upstream of *Vgsc*. In addition to being narrower in genomic distance, the magnitude of the putative sweep is also lower in the Colombian sample than in the Brazilian one. Additionally, one of the genes identified in the prominent sweep on chromosome 2 discussed above is ankyrin gene 2,3/unc44 (AAEL013466); ankyrin proteins are associated with, and directly interact with, voltage-gated sodium channels (Bennett and Baines 2001, Bennett and Chen 2001, Jenkins and Bennett 2001). In *Ae. aegypti*, variation in ankyrin-domain-containing genes has been associated with pyrethroid resistance (Campbell et al. 2019); in addition, the ankyrin repeat region-containing gene AAEL011338 was identified as harboring a locus associated with pyrethroid resistance in a Brazilian sample (Cosme et al. 2022).

Next, we examined two peaks present in the Gabon and Kenya samples. On chromosome 2, a sweep signal present in these two samples stretches from approximately 227.5 Mb to 228.2 Mb, overlapping 9 genes (“*f”, Figure 2, Figure S6f). The two genes with specific functional predictions fully encompassed by this sweep are the mitochondrial ribosomal protein L52 gene AAEL023209, and *SCRB8* (AAEL000227), a class B scavenger receptor. However, just downstream, the Gabon cohort shows another signal of a putative selective sweep, from approximately 229.5 Mb to 231.25 Mb. This peak, which is one of the strongest detected in any cohort, overlaps 39 genes, 19 of which are protein-coding and 10 of which have a specific functional prediction (Table S7). Here, we discuss five known in the literature. *SCRB5* (AAEL011222) has been found to be upregulated in *Ae. aegypti* cells treated with human complement protein hC5a (Giraldo-Calderón et al. 2020). The gene AAEL008073, predicted to bind RNA (VectorBase release 58; (Dickson et al. 2017), is overexpressed in bloodfed *Ae. aegypti* compared to sugar-fed (Bonizzoni et al. 2011); it also displays very low heterozygosity in populations from Mexico, Thailand, and Senegal (Dickson et al. 2017). In addition, dsRNA silencing of this gene and infection with chikungunya in an *Ae. aegypti* cell line results in a significant decrease in chikungunya RNA levels (Dubey et al. 2022). When *notch* (AAEL008069), another nearby gene, is silenced, *in* vitro viral load of DENV-2 (dengue serotype 2) is significantly increased six days post-infection compared to control mosquitoes (Xiao et al. 2014). A splice variant of the gonadotropin-releasing hormone receptor *AAEL011325* is down-regulated in larvae exposed to brackish water compared to freshwater (Durant et al. 2021). Finally, the nicotinic acetylcholine receptor subunit gene *Aaea6* (AAEL008055) is upregulated in response to post-eclosion application of exogeneous juvenile hormone (Zhu et al. 2010), and has also been experimentally identified as the target of the insecticide spinosad (Lan et al. 2021); more generally, nAChR genes are known as the targets of neonicotinoid insecticides (Casida 2018).

Senegal shows a series of putative selective sweeps on chromosome 2 between 312 and 316 Mb (“*g”, Figure 2, Figure S6g). Collectively, these peaks overlap 35 genes, 26 of which are protein-coding and 11 of which have a specific predicted function (Table S7). The aldo-keto reductase AAEL007275 is up-regulated after larval imidacloprid exposure (Riaz et al. 2009), and up-regulated after one generation of selection for resistance to temephos (Saavedra-Rodriguez et al. 2014). The alpha-amylases AAEL010540 and AAEL010532 contain nonsynonymous changes in putative receptors for *Bacillus thuringiensis israelensis* (Bti) Cry toxins that are nearly fixed between control strains and strains selected for resistance to individual Cry toxins (Stalinski et al. 2016). Another study showed AAEL010540 underexpression in strains resistant to certain Cry toxins compared to a control strain (Després et al. 2014). ABC transporter *ABCC13* (AAEL023524) is underexpressed in late dengue infection and overexpressed throughout yellow fever expression *in vivo* (Kumar et al. 2021). The importin alpha gene AAEL012960 is demethylated following infection with a virulent strain of *Wolbachia* (Ye et al. 2013).

In addition, five genes in this region (AAEL012960, AAEL013222, AAEL022016 [alternate gene ID: AAEL013219], AAEL010533, and AAEL023254 [previous gene ID: AAEL013215]) were identified as among the top 25 most differentiated genes between an urban population in Senegal vs. a forest population from Senegal together with a population from Uganda (Crawford et al. 2017); AAEL012960 and AAEL023254 contain exonic SNPs highly differentiated in that comparison. This sweep also overlaps a region identified as containing many variants associated with human specialist ancestry, and as highly diverged based on the Population Branch Statistic, in these specimens (Rose et al. 2020), congruent with our identification of a putative sweep.

Finally, the cohort from Senegal also shows several putative selective sweeps on chromosome 3 between 100 and 140 Mb (“*h”, Figure 2, Fig S6h). The three most prominent peaks in this region overlap 25 protein coding genes, 7 of which have a specific functional prediction (Table S7). The cell division cycle 20 gene (*cdc20*, AAEL004480) is underexpressed after adult females are challenged with either *Escherichia coli* or *Micrococcus luteus* (Choi et al. 2012), and contains a SNP showing significant spatial structuring in populations in Rio de Janeiro (Rašić et al. 2015).

### Selective sweep signatures are enriched for insecticide resistance genes in Colombia

In addition to examining individual sweeps, we looked for categories of genes disproportionately represented in sweeps across the genome. Here, we used the individual SweepFinder2 test sites, distributed approximately every 1 kb throughout the genome. We identified outliers in these test sites at the 99^th^ percentile in each country, with enrichment testing performed in Gowinda with a 1 kb buffer up- and downstream of each gene (Figure S7). We found that eight GO terms were enriched in four populations, with seven (DNA binding; DNA-templated transcription, initiation; chromosome; host cell nucleus; nucleosome; nucleosome assembly; and nucleus) related to DNA or DNA binding. Further examination of the genes underlying this enrichment showed that it was partially attributable to the presence of a cluster of histone genes at approximately 340 Mb on chromosome 3, overlapped by outlier windows in the four populations (Figure S8). This highlighted the possibility of clusters of genes influencing the enrichment results, so we implemented a permutation test to test for significantly enriched GO terms after accounting for physical location (see Methods). After controlling for gene clustering this way, no GO terms were significantly enriched at a *q* value of 0.05 when examining genes overlapping outliers windows at the 99^th^ percentile. These results highlight the need to account for physical clustering of genes.

We also tested whether genes potentially associated with insecticide resistance genes were enriched in outlier windows. To do this, we assembled a list of 152 cytochrome P450 genes, glutathione transferase genes, and carboxylesterase genes by searching the annotated gene file available through VectorBase (release 54). We then manually added AAEL023266 (*Vgsc*), and AAEL023844 (a carboxylesterase-like gene that is part of a cluster of *CCE* genes and has been extensively associated with insecticide resistance [(Cattel, Haberkorn, et al. 2021, Cattel, Minier, et al. 2021, Sene et al. 2021)])—we note that the decision to include *Vgsc* in enrichment testing was made prior to conducting our SweepFinder2 scan. Outlier windows at the 99^th^ percentile were significantly enriched for these genes in Colombia, but not the other five countries (however, the *p*-value for enrichment in the Brazil cohort was 0.05).

### Strong but variable signatures of selection around Vgsc in South America

Given its important role in resistance to insecticides in numerous species, we conducted a more thorough analysis of *Vgsc* and the surrounding region. We began by conducting principal component analysis of SNPs inside *Vgsc* (from 315,926,360 to 316,405,639 on chromosome 3), which showed specimens clustering into an approximate triangle (Figure 3a). The point of the triangle is anchored on the right side by a cluster of Senegalese, Gabonese, and Kenyan specimens, and the *x-*axis (the first principal component, which explains 25% of the variance), approximately divides African specimens from American specimens, with, however, much intermixing. The base of the triangle, on the left, comprises American specimens only; the *y*-axis (the second principal component, explaining 15% of the variance) shows American specimens in three distinct and roughly equidistant clusters, with some spread between them. All Colombian and Brazilian specimens can be easily assigned to one of the clusters, and a subset of the specimens from the USA take intermediate positions.

**Figure 3.**
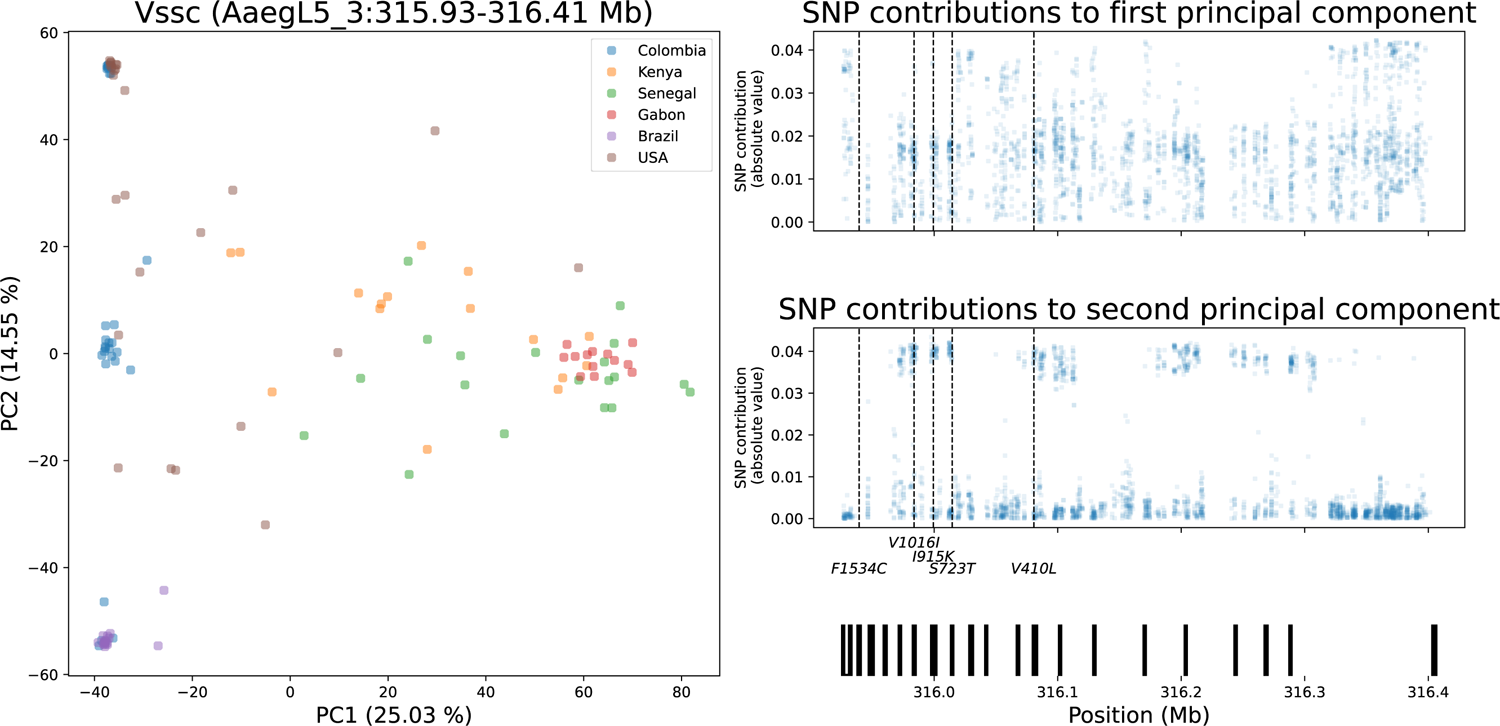
Principal component analysis and loadings of *Vgsc*. Vertical dashed lines on the loadings panels indicate the locations of the five focal loci listed below the plot. The vertical black bars at the bottom right show the location of exons in *Vgsc*.

We then investigated genotypes for each specimen at a number of non-synonymous SNPs previously shown to be either causally linked or associated with pyrethroid resistance (our “focal sites”): F1534C (Kawada et al. 2009, Harris et al. 2010, Hu et al. 2011, Du et al. 2013, Hirata et al. 2014), V1016I (Saavedra-Rodriguez et al. 2007, Saavedra-Rodriguez et al. 2008, Harris et al. 2010, Du et al. 2013), S723T (Saavedra-Rodriguez et al. 2019, Mack et al. 2021), V410L (Haddi et al. 2017, Saavedra-Rodriguez et al. 2019), I915K (Mack et al. 2021), and Q1853R (Kelly et al. 2021) (Table 1; we use the *Musca domestica* nomenclature). F1534C was not called in any specimen in our filtered data set, although we were later able to genotype this locus as discussed below (these genotypes were not included in the PCA but do appear in Table 1); Q1853R was confidently called as invariant in 103 specimens and either called as invariant with low confidence or uncalled in the remaining 15, and thus excluded from further analysis. The remaining four sites were confidently called and were polymorphic in our data, with high LD between them in the Colombian sample. The genotypes at these four positions also correspond to the three clusters seen on the left side of Figure 3a, with the bottom cluster corresponding to the specimens homozygous for the reference or wild-type alleles at the four sites, the top cluster corresponding to the specimens homozygous for alleles associated with resistance, and the middle group corresponding to specimens heterozygous at all four sites.

**Table 1.**
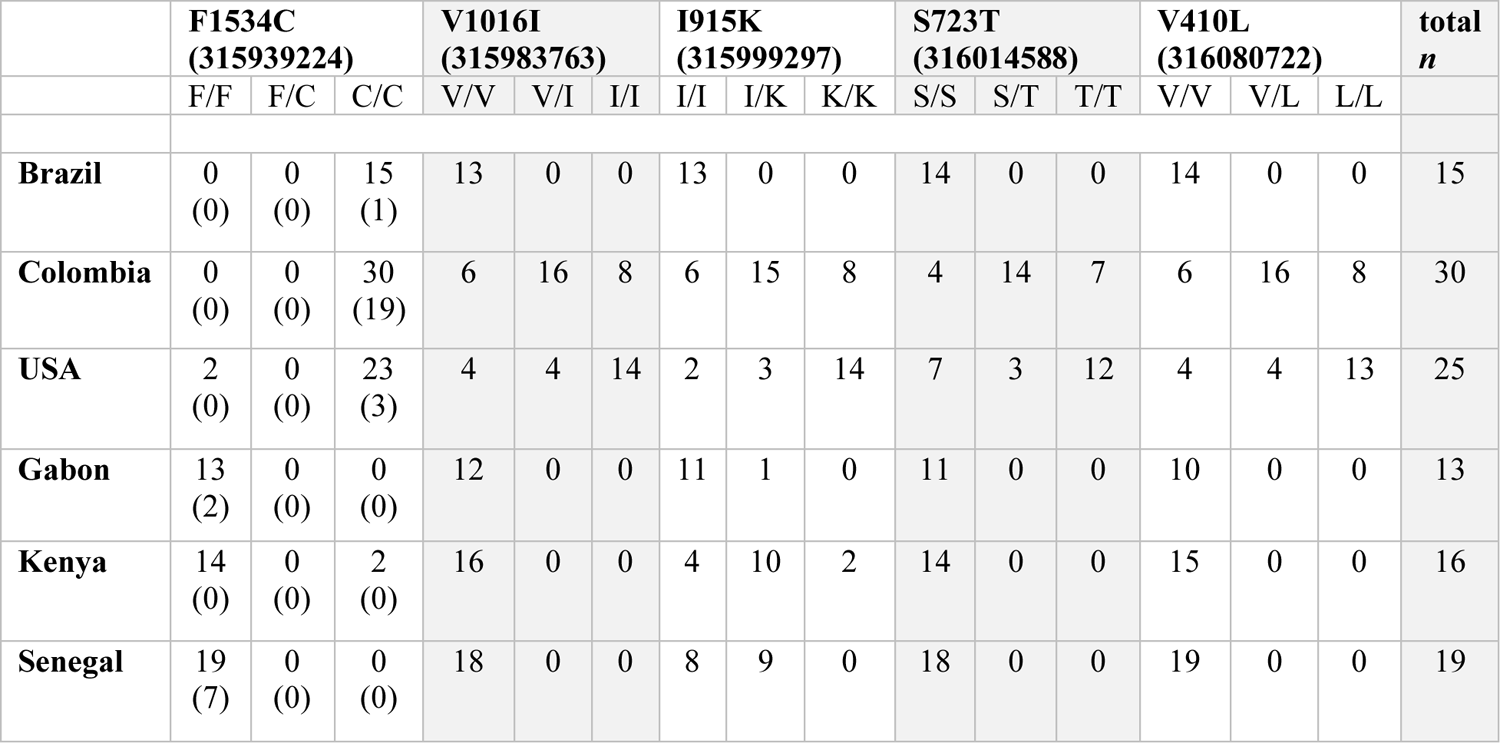
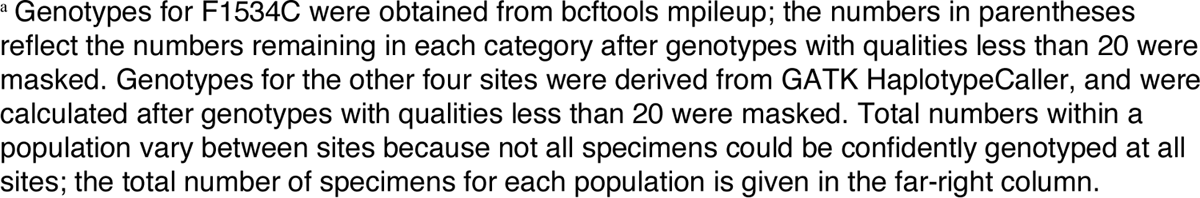
Genotypes at selected loci previously identified as associated with pyrethroid resistance in *Vgsc*.^a^

|  | **F1534C**<br>**(315939224)** |  |  | **V1016I**<br>**(315983763)** |  |  | **I915K**<br>**(315999297)** |  |  | **S723T**<br>**(316014588)** |  |  | **V410L**<br>**(316080722)** |  |  | **total**<br>***n*** |
| --- | --- | --- | --- | --- | --- | --- | --- | --- | --- | --- | --- | --- | --- | --- | --- | --- |
|  | F/F | F/C | C/C | V/V | V/I | I/I | I/I | I/K | K/K | S/S | S/T | T/T | V/V | V/L | L/L |  |
| **Brazil** | 0<br>(0) | 0<br>(0) | 15<br>(1) | 13 | 0 | 0 | 13 | 0 | 0 | 14 | 0 | 0 | 14 | 0 | 0 | 15 |
| **Colombia** | 0<br>(0) | 0<br>(0) | 30<br>(19) | 6 | 16 | 8 | 6 | 15 | 8 | 4 | 14 | 7 | 6 | 16 | 8 | 30 |
| **USA** | 2<br>(0) | 0<br>(0) | 23<br>(3) | 4 | 4 | 14 | 2 | 3 | 14 | 7 | 3 | 12 | 4 | 4 | 13 | 25 |
| **Gabon** | 13<br>(2) | 0<br>(0) | 0<br>(0) | 12 | 0 | 0 | 11 | 1 | 0 | 11 | 0 | 0 | 10 | 0 | 0 | 13 |
| **Kenya** | 14<br>(0) | 0<br>(0) | 2<br>(0) | 16 | 0 | 0 | 4 | 10 | 2 | 14 | 0 | 0 | 15 | 0 | 0 | 16 |
| **Senegal** | 19<br>(7) | 0<br>(0) | 0<br>(0) | 18 | 0 | 0 | 8 | 9 | 0 | 18 | 0 | 0 | 19 | 0 | 0 | 19 |

We also looked at the contribution of each SNP to the groupings in the PCA (Figure 3b), and specifically to the second principal component, along which many American specimens cluster into discrete groups. The loadings for this principal component show several clusters of SNPs with some of the highest contributions in the PCA. When we plot the location of five associated resistance SNPs, we see that, except for F1534C (which was not included in the PCA, as previously described), they correlate with clusters of SNPs that are outliers in terms of their contribution to the second principal component. In addition to the clusters associated with the known resistance loci, there are additional clusters in the proximal portion of the gene. We identified three SNPs (316268746, 316268894, and 316269032) with a PCA loading of greater than 0.03 that overlap two adjacent exons; all three SNPs are found in the untranslated region. *Vgsc* is extensively alternatively spliced (Lin et al. 2009), so these SNPs may be relevant in that regard, or may influence expression levels; however, they may also have no functional importance and simply be in high LD with the focal polymorphisms and/or another causal polymorphism not present in our filtered data set.

Examining nucleotide diversity in the region of *Vgsc*, from 310 to 320 Mb on chromosome 3, we observe a decrease in diversity in the South American populations from approximately 315 Mb to 318 Mb (Figure 4a). The specimens from the USA do not show such a drop; however, this heterogeneous cohort includes specimens with ancestry from multiple source populations (Lee et al. 2019, Kelly et al. 2021), which could give the appearance of high nucleotide diversity even if diversity is low within some ancestry groups. Relative to the Brazilian sample, the nucleotide diversity of the Colombian sample is elevated within *Vgsc* itself, and in the distal region to approximately 318 Mb, where nucleotide diversity increases in the Brazilian sample. This is in keeping with the number of clusters observed for each cohort in the PCA of the *Vgsc* region as well as the genotypes shown in Table 1: one cluster for Brazil, and three for Colombia, indicating multiple haplotypes in the latter but not the former.

**Figure 4.**
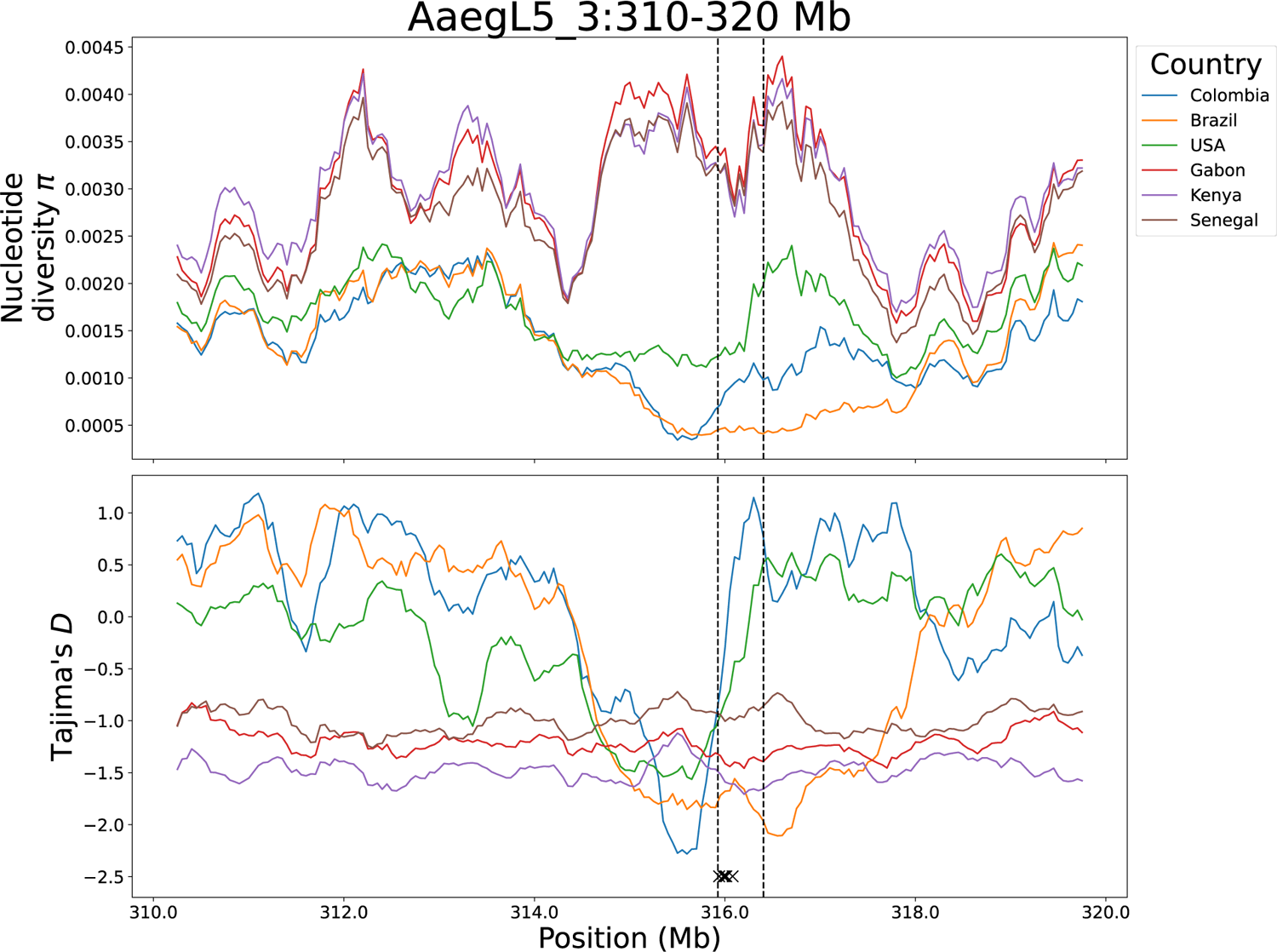
Nucleotide diversity and Tajima’s *D* in six cohorts, on chromosome 3 from 310 to 320 Mb. The vertical dashed lines mark the boundaries of *Vgsc* (315,926,360-316,405,639). The five Xs at the bottom of the Tajima’s *D* plot indicate the locations of the five focal loci.

Although the SweepFinder2 results suggest a strong skew in allele frequencies in this region, to more directly examine this, we calculated Tajima’s *D*. In the American samples, Tajima’s *D*, which is otherwise generally higher than in the African samples, drops sharply proximal to *Vgsc* (Figure 4b). This drop is especially pronounced in the Colombian sample. However, within *Vgsc* itself, Tajima’s *D* is sharply elevated; this is in keeping with the presence of multiple haplotypes in this region within Colombia.

Finally, linkage disequilibrium (*r*^2^, calculated as described in the Methods) between individual variants within *Vgsc* in the three populations from the Americas shows that regions of elevated LD are particularly prominent in the South American populations (Figure 5). One such block occurs from approximately 315.97 Mb to 316.08 Mb—we note that this block contains the V1016I, I915K, S723T, and V410L polymorphisms, which are all in high LD with one another in the Colombian sample (Table 2; compare to USA sample, Table S8). This region also shows noticeably elevated LD with the region from 316.31 Mb to the end of the gene. In addition, another block of elevated LD occurs from approximately 316.17 Mb to 316.30 Mb. The USA population, in contrast, shows a more diffuse pattern of elevated LD, spread across the gene.

**Figure 5.**
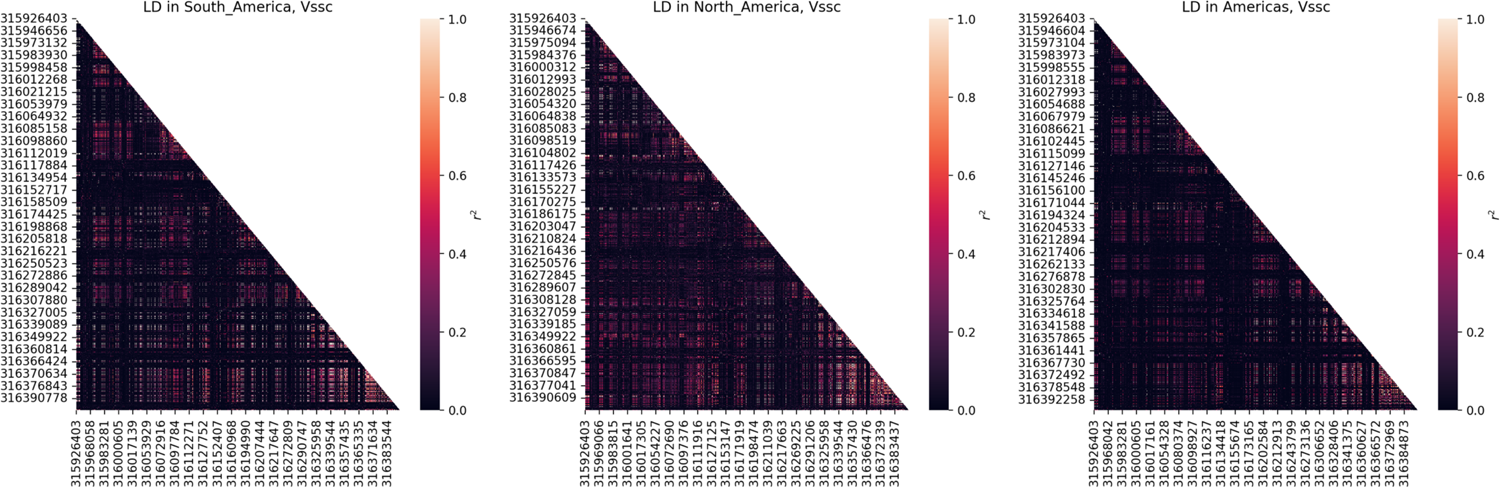
Linkage disequilibrium across *Vgsc* in the North American cohort (i.e. USA), the South American cohorts (Brazil and Colombia combined), and the combined American cohort (Brazil, Colombia, and USA combined).

**Table 2.**
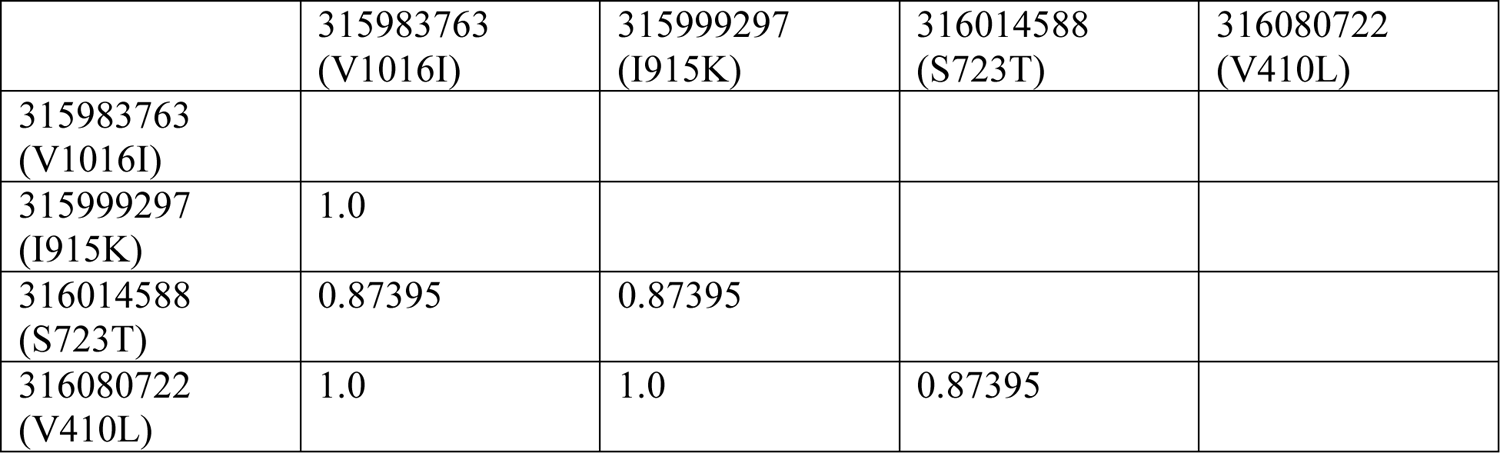
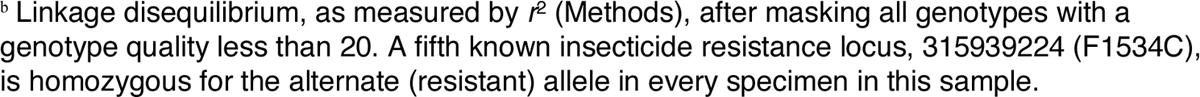
Linkage disequilibrium between four known or putative insecticide resistance loci in the Colombian sample.^b^

|  | 315983763<br>(V1016I) | 315999297<br>(I915K) | 316014588<br>(S723T) | 316080722<br>(V410L) |
| --- | --- | --- | --- | --- |
| 315983763<br>(V1016I) |  |  |  |  |
| 315999297<br>(I915K) | 1.0 |  |  |  |
| 316014588<br>(S723T) | 0.87395 | 0.87395 |  |  |
| 316080722<br>(V410L) | 1.0 | 1.0 | 0.87395 |  |

<sup>b</sup> Linkage disequilibrium, as measured by $r^2$ (Methods), after masking all genotypes with a genotype quality less than 20. A fifth known insecticide resistance locus, 315939224 (F1534C), is homozygous for the alternate (resistant) allele in every specimen in this sample.

These findings are particularly striking because the Brazilian sample, though wild-type for all previously identified SNPs associated with pyrethroid resistance that could be confidently genotyped in our data, still shows a sharp drop in nucleotide diversity and Tajima’s *D* characteristic of a selective sweep; in addition, SweepFinder2 identified a strong candidate selective sweep in both the Brazilian and Colombian samples in this region. Indeed, SweepFinder2’s CLR peak was substantially higher and broader in Brazil than Colombia. These findings, as well as the well-known role of the F1534C polymorphism in insecticide resistance, led us to re-examine this locus in our data. F1534C was also not present in the unfiltered variants, indicating this site had not been simply filtered out. In our set of unfiltered variants, we found a total lack of coverage in all specimens in the vicinity of this SNP, spanning from approximately 315,937,705 bp to 315,945,225 bp. However, coverage in the original alignments appeared relatively normal (Figure S9a). Inspection of the alignments revealed low mapping quality in this region (Figure S9b), with a high proportion of reads aligning to this region also mapping to another region, scaffold NIGP01000811; reads that align equally well to multiple locations are given a mapping quality of 0 (Li et al. 2008) and thus are not considered by GATK’s HaplotypeCaller using the default settings, which remove all reads with a mapping quality below 10 (https://gatk.broadinstitute.org/hc/en-us/articles/360036889192-MappingQualityReadFilter). The similarities between this scaffold and *Vgsc* have previously been noted (Itokawa et al. 2019). Aligning NIGP01000811 to chromosome 3 of AaegL5 revealed that most of the scaffold aligns to the vicinity of the *Vgsc* gene, but the first approximately 4.9 kb do not confidently align anywhere on chromosome 3 (Figure S10).

We therefore re-genotyped chromosome 3 from 315 to 317 Mb, as well as the entirety of NIGP01000811, using bcftools version 1.9 mpileup, which has a default minimum mapping quality of 0 (https://samtools.github.io/bcftools/bcftools.html). Using this method, we called putative genotypes at F1534C, revealing that every South American specimen is homozygous for the resistant allele (Table 1). In the Brazilian sample in particular, the genotypes are of very low quality, reflecting low confidence in the genotype calls. However, this is expected, as read mapping qualities influence the calculation of genotype likelihoods and therefore qualities (Danecek et al. 2021).

While one explanation for this apparent duplication is a technical artifact of assembly, we cannot rule out the presence of a genuine duplication, and therefore we are less confident in the genotypes at F1534C than at the other putative insecticide resistance loci; however, we present these data given the established importance of this SNP on insecticide resistance and vector control. To assess the effect of using a different variant caller on clustering observed in this region, we repeated the principal components analysis using the variants genotyped by bcftools (Figure S11); the triangular pattern observed in the calls generated by GATK is generally recapitulated, though excluding low-quality genotypes introduces a great deal of noise.

In addition, we calculated nucleotide diversity separately within two sets of South American samples: those in the top and bottom clusters of the *Vgsc* PCA (Figure S12). This reveals a sharp valley in nucleotide diversity in the bottom cluster, which contains the specimens carrying a known resistance allele at F1534C only, and an even sharper decrease in nucleotide diversity in the top cluster, which contains specimens carrying known or putative resistance alleles at all five focal loci.

Finally, drawing firm conclusions about the North American sample published in Lee et al. (2019) (Lee et al. 2019) and analyzed here is difficult because of the multiple founding populations and small per-cluster sample sizes; however, when we decompose the portion of the sample originating in California into the clusters noted in Lee et al. (two clusters from central California, and one cluster from southern California plus one site in central California), we see decreased nucleotide diversity upstream of *Vgsc* and increased nucleotide diversity within *Vgsc* itself in two of the three clusters (Figure S13), consistent with the data observed in Colombia. In addition, Mack et al. (2021) (Mack et al. 2021) observed high frequencies of the resistant allele at the five focal loci studied here in several localities in central California; data from Kelly et al. (2021) (Kelly et al. 2021) similarly suggest an increase in the frequency of resistance alleles in Californian samples between 2014 and 2018. Finally, when examining the PCA of high-quality SNPs within *Vgsc*, a subset of the USA cohort clusters very closely with the Colombian specimens that carry all five focal insecticide resistance alleles (Figure 3). Thus, it appears likely that the resistance haplotype observed in Colombia is present in parts of North America as well.

## DISCUSSION

These whole-genome data from six country-level cohorts of *Aedes aegypti* on three continents allowed us to consider the history of this species in detail. In general, our findings are consistent with the known history of this species. Tajima’s *D* values are negative in all three African cohorts, while they are either positive, or very close to zero, in the three American cohorts; in addition, the Senegal population shows values of Tajima’s *D* intermediate between the Gabon and Kenyan cohorts, and the American cohorts. This is in keeping with previous studies of populations from these regions (Crawford et al. 2017), and is consistent with the idea of a population bottleneck associated with domestication, perhaps followed by a second bottleneck associated with expansion out of Africa (Crawford et al. 2017). The positive Tajima’s *D* in American cohorts may reflect a relatively recent contraction due to the continent-wide eradication effort, but population genetic evidence of subsequent recovery and expansion will require larger sample sizes to detect the excess of very rare alleles produced by such a change (Tennessen et al. 2012).

Of the three American populations, Tajima’s *D* is highest in the Brazilian sample; linkage disequilibrium values are also sharply elevated in this population compared to all others. Eradication of *Ae. aegypti* was claimed to be complete in Brazil (Soper 1965), but no such claim was made for Colombia. Our observations are consistent with a historical reduction in population size in Brazil, perhaps due to a founder effect during re-infestation from another country. The observation of two specimens from Brazil and Colombia, respectively, showing kinship consistent with a third-degree relationship is also consistent with previous observations of clustering between Brazilian and Colombian (Kotsakiozi et al. 2017) populations or other populations from Central and northern South America (Monteiro et al. 2014). The USA cohort has lower Tajima’s *D* values, and lower linkage disequilibrium values, than the other two cohorts from the Americas; however, this sample is heterogeneous (Lee et al. 2019), so these results should be interpreted cautiously.

In addition, we present the first genome-wide scans for positive selection in *Aedes* using SNP-resolution data, identifying candidate selective sweeps in six countries. In multiple cohorts, we identify signatures of strong selective sweeps, often with CLR scores well over 1000, near loci implicated in insecticide resistance, including the voltage-gated sodium channel gene *Vgsc* (Brazil, Colombia); glutathione s-transferases (Brazil, Colombia); a cytochrome P450 (Brazil); and a nicotinic acetylcholine receptor subunit gene (Gabon).

We caution that genes associated with insecticide resistance may be disproportionately well-annotated, and are likely to be disproportionately well-investigated, because of their obvious public health implications. Some sweep signatures overlap a large number of genes, many of which whose functions are currently unannotated, making detailed conclusions about the targets of selection impossible; in addition, the peak of a sweep signature may not always correspond to the target of selection (Schrider et al. 2015). However, insecticide usage is known to cause strong selective pressure in many other insects (Garud et al. 2015, Miles et al. 2017, Walsh et al. 2018, Weedall et al. 2020, Xue et al. 2021, Pélissié et al. 2022), and the variety of genes implicated in insecticide resistance occurring near these sweeps is a potential sign of *Ae. aegypti*’s agility in responding to these insecticides.

The putative selective sweep that is perhaps the most interesting, because of the contrasting patterns shown in two different South American countries, is at *Vgsc*. Examining this genomic region in Colombia reveals two observations: one, somewhat unexpectedly, the putative selective sweep in this population appears to be localized upstream of the gene, not within *Vgsc* itself, based on results from SweepFinder2 as well as elevated Tajima’s *D* within *Vgsc*. Two, the clustering of the Colombian sample into three tight groups along principal components suggests that two long-range haplotypes linking several loci associated with insecticide resistance are present in the population, with one haplotype containing the resistant alleles at at least four previously identified insecticide resistance SNPs, and the other containing the wild-type alleles. Indeed, examining the PCA loadings shows that these polymorphisms line up very well with clusters of SNPs driving the separation between PCA clusters. We note that this pattern of three equidistant clusters on a PCA is the characteristic signature of a polymorphic chromosomal inversion (Ma and Amos 2012, Love et al. 2019); however, chromosomal inversions are difficult to detect *de novo* from short read data, and long-read data are likely necessary to either confirm or rule out the possibility of an inversion in this region.

Examining this same region in the Brazilian sample reveals a very different picture. Here, the putative selective sweep is centered around *Vgsc*; however, PCA results, as well as genotypes at individual loci associated with resistance, suggest that the haplotype of resistant alleles is absent in this population. Indeed, all the Brazilian specimens carry the wild-type susceptible alleles at four of the five insecticide resistance loci examined. At the fifth loci, F1534C, a partial duplication of the *Vgsc* gene in the assembly prevents confident genotyping; however, if we assume that this duplication is technical in origin and that reads mapping to the duplicated region should instead be assigned to *Vgsc* itself, we find that the Brazilian sample is completely homozygous for the resistant allele at this locus (Table 1). This suggests that the strong selective sweep observed in this population is driven either by F1534C, by other loci that have yet to be identified, or both, but not by V410L, S723T, I915K, or V1016I, all of which are absent from our Brazilian cohort but present in the adjacent country of Colombia.

We hypothesize that the Brazilian and Colombian samples may in fact have been subject to the same selective sweep at this locus, despite the absence of the signal of such a sweep in *Vgsc* itself in the Colombian sample. Our proposed explanation to this apparent discrepancy involves the putative long-range haplotypes present in the Colombian population. The presence of two such haplotypes, both at intermediate frequency, would necessarily skew the SFS toward intermediate-frequency alleles, thereby increasing Tajima’s *D* in this region. In effect, we suggest that in Colombia, a second, and ongoing, selective sweep confined to only part of the *Vgsc* gene, has partially eroded the signal of an earlier sweep that encompassed the entire gene and some of the flanking regions. Alternatively, it could be that this haplotype is being maintained by some form of balancing selection in the Colombian sample, or its intermediate frequency has resulted from an influx of migrant haplotypes from another population, rather than ongoing positive selection. However, the high LD between V410L, S723T, I915K, or V1016I, as well as the lack of diversity among both the resistant and wild-type backgrounds (Fig S11), seem to contradict these alternative explanations; we also note that V410L and V1016I have previously been noted to be in LD with each other in other locations (Vera-Maloof et al. 2015, Saavedra-Rodriguez et al. 2018, Ayres et al. 2020, Baltzegar et al. 2021). If our hypothesis is correct, the frequency of the haplotype bearing these resistance mutations may continue to increase in the coming years under continuing selective pressure from insecticide use. Indeed, the Brazilian sample is from 2012, so allele frequencies at these loci may be very different in the present day, as the resistant alleles at V1016I and F1534C are common (Linss et al. 2014, David et al. 2018, Melo Costa et al. 2020, Cosme et al. 2022) and increasing in frequency (Linss et al. 2014) in some parts of Brazil. This situation highlights the need for an increased use of temporal data in vector genomics.

An apparent sweep in the vicinity of *Vgsc* has been observed previously in a sample from southern Mexico (Saavedra-Rodriguez et al. 2019); in addition, data from North America must be interpreted with caution for reasons previously discussed, but are consistent with reduced diversity in the vicinity of *Vgsc* and an increase in insecticide resistance alleles over time. Taken all together, these data suggest this long-range haplotype may not be limited to Colombia, and could be spreading more widely across the Americas.

An earlier origin of F1534C has been posited before (Saavedra-Rodriguez et al. 2018), and would be consistent with previous observations that F1534C is more commonly found in the absence of V410L (Haddi et al. 2017, Melo Costa et al. 2020) and V1016I (Linss et al. 2014, Vera-Maloof et al. 2015, Melo Costa et al. 2020) than vice versa; in addition, in Iquitos, Peru, F1534C was directly observed to reach near-fixation before V1016I became prevalent (Baltzegar et al. 2021). These findings also lead us to speculate about the catalyst for this putative earlier sweep. Pyrethroids and DDT both target the voltage-gated sodium channel, and F1534C is known to confer resistance to DDT as well as to type I pyrethroids (Harris et al. 2010, Hu et al. 2011, Du et al. 2013); cross-resistance to both insecticides has been observed in *Ae. aegypti* (Chadwick et al. 1977). Populations of *Ae. aegypti* across Colombia are still highly resistant to DDT (Fonseca-González et al. 2011), though its use for vector control has been banned in that country since 1993 (Fonseca-González et al. 2011). Though the F1534C allele is present at high frequency in the USA sample and a subset of the sample shows reduced nucleotide diversity in this region, we do not see a putative selective sweep at this locus at anywhere near the magnitude of the other American populations. Although it is possible that heterogeneity within the USA sample may be somewhat masking the signature of a strong sweep, it is also possible that that the selective pressure on the sweeping allele (whether it was F1534C or otherwise), in populations in the USA, may not have been strong enough to drive one haplotype to high frequency as quickly as in South America. Unlike in South and Central America, continent-wide eradication of *Ae. aegypti* was never attempted in North America. While we can only speculate that the apparent sweep of F1534C in South America may be tied to the attempted eradication with DDT, it is worth noting that DDT, not pyrethroids, are likely to have presented the strongest anthropogenic selection pressure these populations have ever experienced.

## Conclusions

Here we describe a dataset of 30 *Aedes aegypti* genomes collected from Colombia. We combined these data with whole genomes from five other countries from Africa and the Americas to examine the genomic impacts of population structure, potential past demographic events, and natural selection on this important vector species. While this analysis has revealed some clues about potential recent demographic events (e.g. domestication-associated bottlenecks, and recent migrations), future work with broader sampling should test these possibilities explicitly. Perhaps our clearest contribution is the strong signatures of positive selection that we identify around numerous insecticide resistance genes. These results paint a compelling picture of strong selective pressures, and a population that appears to be able to rapidly adapt to these pressures in part due to a fairly large number of genes that can potentially confer resistance phenotypes; these results are consistent with previous observations that a variety of loci and pathways underlie *Ae. aegypti*’s response to insecticides in general and pyrethroids in particular (Saavedra-Rodriguez et al. 2008, Saavedra-Rodriguez et al. 2012, Paris et al. 2013, Saavedra-Rodriguez et al. 2014). We also note that because *Aedes agypti* has a large and expanding global population in terms of its census size, adaptation may not be mutation-limited even at individual sites. For example, two different resistance mutations at the same codon in *Vgsc*, V1016G and V1016I, are prevalent in Asia and the Americas, respectively, and at least one mutation in *Vgsc*, F1534C, appears to have occurred recurrently (Cosme et al. 2020). Moreover, the ability to occasionally travel long distances through the aid of human transportation may allow for the sharing of resistance alleles across populations. Thus, we argue that continued efforts to identify selected alleles in *Aedes aegypti* populations and to monitor their spread will be vital for informing vector control efforts.

## METHODS

### Sample collection

We sampled two urban locations in Medellín (Comuna 16 Belen, Comuna 13 San Javier) and in a rural locality near the nature reserve area of Río Claro in the department of Antioquia. Medellín has an average altitude of 1,500 meters, and a mean annual temperature of 22.5°C. Río Claro has an average altitude of 1,750 meters, and a mean annual temperature of 22°C. We also collected samples from Cali, capital of Valle del Cauca department (3.3817 N, −76.55237 W). Cali has an average altitude of 1,479 meters, average temperature of 24 °C, and average annual precipitation of ∼1,050 mm. The Cañón del Río Claro Nature Reserve is located between the municipalities of Sonsón, Puerto Triunfo and San Luis, to the southwest of the department of Antioquia (5.90 N, −74.86 W; (Vivero et al. 2010)).

Our collection focused on *Ae. aegypti* larval stages, and their main potential breeding grounds (pools and tanks used to store drinking water) were inspected. Larvae were then transported in Whirl-Pak plastic bags to the insectary of the Program for the Study and Control of Tropical Diseases (PECET) of the University of Antioquia, Medellín, Colombia (27 ± 2 ° C; relative humidity 80 ± 10%; photoperiod: 12 light hours), where stages III and IV were individualized and their breeding continued for entomological series, reference collection and molecular processing of adults. Larvae, pupae and adults were sorted following the protocols suggested by the Biosystematics Unit, Division of Entomology, Walter Reed 1997. To assign to species (*Ae. aegypti*), we used external morphological characters of larvae, pupae and adult females.

### Ethics statement

All specimen collections were performed following the guidelines of the Colombian decree N° 1376. No specific permits were required for this study. Mosquitoes were collected on private property and permission was received from landowners before sampling.

### DNA extraction and sequencing

We extracted genomic DNA from individual wild-caught adult *Aedes* in Colombia using Genetra Puregene Tissue Kits (Qiagen, Valencia, CA, USA), constructed barcoded libraries for sequencing using KAPA HyperPrep kits (Roche Sequencing, Pleasanton, CA) with a target fragment size of 500 bp. We assessed the size of the library inserts using a Tapestation 4150 (Agilent, Santa Clara, CA) and dsDNA screentape (Agilent, #5067-5365). Finally, we pooled the samples and sequenced them in a NovaSeq6000 S2 (Illumina) machine, generating 2×150bp reads. The target coverage was 15X per sample. This effort yielded 40 sequenced genomes.

To these specimens we added publicly available sequence data (Lee et al. 2019, Rose et al. 2020), as previously noted.

### Quality control, alignment, and variant genotyping

We used FASTQC 0.11.9 (Andrews et al. 2010) to check initial read quality, and Trimmomatic 0.39 (Bolger et al. 2014) to remove adapters and trim regions of low quality at the beginnings and ends of reads. After trimming, we re-checked read quality using FASTQC.

For newly sequenced specimens, read group information was extracted using bash scripting. Reads were aligned to AaegL5, retrieved from VectorBase (Matthews et al. 2018, Giraldo-Calderón et al. 2022) using the mem tool of bwa-mem2 version 2.1 (Vasimuddin et al. 2019), converted to BAM format, and, for specimens split across multiple lanes, merged across lanes using samtools version 1.11 (Danecek et al. 2021).

We used MarkDuplicatesSpark in GATK version 4.1.9.0 (DePristo et al. 2011, Poplin et al. 2018) to mark optical duplicates and sort reads. BAM files were then indexed with samtools index. Quality of the resulting BAM files was assessed with qualimap version 2.2.2d (Okonechnikov et al. 2016) bamqc and with picard CollectMultipleMetrics.

Sites were genotyped with GATK HaplotypeCaller (Poplin et al. 2018) independently for each specimen, parallelized across chromosomes, using the EMIT_ALL_CONFIDENT_SITES flag. The resulting gVCF files were merged with picard GatherVcfs and indexed with GATK IndexFeatureFile. Quality metrics for each specimen were compiled from FASTQC, picard, and qualimap using multiqc version 1.9 (Ewels et al. 2016).

Genomic VCF files for each specimen were then turned into batch variant calls using GATK. Specifically, specimens were compiled using GenomicsDBImport and genotyped using GenotypeGVCFs with the include-non-variant-sites flag, parallelized across the genome as necessary for efficiency. The resulting VCFs were merged using picard GatherVcfs and filtered to SNPs only using the view command from bcftools 1.8 and 1.9 (Danecek et al. 2021). The quality control and alignment through individual variant calling steps, and the batch genotyping steps, were implemented as two nextflow v. 20.10.0 (Di Tommaso et al. 2017) workflows, available on GitHub (https://github.com/rrlove/aedes_pipeline).

The merged VCFs were converted into zarr format using scikit-allel 1.3.2 (DOI:10.5281/zenodo.4759368). Variants were filtered in python using scikit-allel. For our focal sample from Colombia, we removed specimens with a mean coverage below 15x or fewer than 70% of reads mapping to the reference genome. In the combined sample, we also removed invariant sites, and sites with indels. Based on GATK best practices for hard filtering (https://gatk.broadinstitute.org/hc/en-us/articles/360035890471-Hard-filtering-germline-short-variants), the following six hard filtering criteria were implemented, with sites required to pass five of the six filters: 1) variant quality by depth (QD) greater than or equal to 2; 2) variant strand bias as estimated by Fisher’s exact test (FS) less than 40; 3) variant strand bias as estimated by the symmetric odds ratio test (SOR) less than 4; 4) mapping quality (MQ) greater than or equal to 40; 5) mapping quality rank sum test (MQRS) between −5 and 5, inclusive; 6) site position within reads rank sum test (RPRS) between −3 and 3, inclusive. In addition, sites were removed if less than 2/3 of specimens were called with a genotype quality of 20 or greater.

To identify and remove closely related specimens from our sample, we calculated relatedness using KING-robust (Manichaikul et al. 2010) with the –kinship flag for all three chromosomes. To remove any possible interference of the *m* locus and surrounding region (Fontaine et al. 2017, Matthews et al. 2018) with inferred kinship coefficients, we repeated this analysis using chromosomes 2 and 3, only. Following (Rose et al. 2020), we used a kinship coefficient cutoff of 0.20 in American specimens and 0.05 in African specimens, to account for higher levels of inbreeding found in American specimens. If a pair of specimens had a kinship greater than this threshold, we removed the specimen with lower coverage.

### Patterns of genetic diversity within and between populations

We visualized genetic structure in our sample using principal components analysis on each chromosome and on the three chromosomes together. SNPs were excluded if their minor allele frequency was 5% or less in the total sample. Genotypes were converted to the number of alternate genotypes present (0 for homozygous reference, 1 for heterozygous, 2 for homozygous alternate), and principal component analysis was done using scikit-allel. Plots were visualized using the first two principal components.

We also examined patterns of genetic diversity and divergence across the genome. Nucleotide diversity was calculated in nonoverlapping 500 kb windows for each of the six countries using the windowed_diversity function in scikit-allel. Tajima’s *D* was calculated similarly, using the windowed_tajima_d function. For visualization, both statistics were also calculated in sliding 5 Mb windows with a step size of 500 kb. *F_ST_* between each pair of countries was calculated using scikit-allel’s moving_weir_cockerham_fst function, using nonoverlapping windows of 500,000 variants. For computational tractability, linkage disequilibrium (LD) was calculated in parallel in 50 Mb genomic regions (or one smaller segment at the end of each chromosome). In each region, linkage disequilibrium was calculated within each cohort in 500 kb nonoverlapping windows. Linkage disequilibrium was summarized by *r*^2^ as calculated using scikit-allel’s windowed_r_squared function, which uses the method of Rogers and Huff (Rogers and Huff 2009), which does not require phasing. Before calculations, genotypes with a genotype quality below 20 were masked; then, non-segregating variants were removed (these might occur because a position is invariant, or is masked due to low genotype quality, in the subset of specimens selected for analysis).

### Scanning for selective sweeps

Putative selective sweeps were identified in each cohort using SweepFinder2 (DeGiorgio et al. 2016), using the empirical site frequency spectrum (SFS) generated from all SNPs segregating in the focal cohort as the background SFS, as well as a genetic map (Matthews et al. 2018). For computational tractability, SweepFinder2 was run in chunks of 50 Mb (or, at the end of each chromosome, one segment containing the remainder), with a grid size of 1000, representing a test site approximately every kilobase. On each chromosome, the 50 Mb chunks overlapped by 5 Mb, and results were plotted to visually check for edge effects before merging. Finally, results for each test site were averaged within 100 kb and 10 kb nonoverlapping windows, for visualization at different scales.

### Enrichment analysis

We used functional enrichment analysis with Gowinda version 1.12 (Kofler and Schlötterer 2012) to look for overrepresented Gene Ontology (GO) terms annotated to genes overlapping putative selective sweeps identified by SweepFinder2. First, the loci (here a “locus” refers to an individual test site, spaced approximately every 1 kb) with sweep likelihoods in the top 1% of all loci were identified. These loci were used as the candidate site file for Gowinda, with all loci queried by sweepfinder, rounded to the nearest base pair, used as the total site file. We included 1,000 bases up- and downstream from each gene to account for possible sweeps in regulatory regions. For each cutoff we ran 1,000,000 simulations, using the “gene” mode, with the following parameters: —min-significance 1 —min-genes 1 —gene-definition gene —gene-definition updownstream 1000. The gene set file was obtained by collapsing entries in the VectorBase *Ae. aegypti* version 54 gene annotation file by GO term.

Many of the genes in the gene sets with the lowest *q*-values were found in clusters, suggesting the potential for physical proximity and linkage disequilibrium to influence these results. Therefore, we implemented an additional enrichment test (https://github.com/SchriderLab/permEnrichmentTest) based on the approach of permuting the association between scores and tested loci. Specifically, this relationship was permuted by joining the three chromosomes into an artificial circular genome, then “rotating” all scores around the circle by a random interval. By preserving relationships between adjacent loci, this approach offers the ability to control for linkage. As before, we ran this enrichment analysis using the top 1% of all tested sites. For each cutoff, we ran 10,000 permutations of the SweepFinder2 scores before asking which GO terms were enriched in the sweep set relative to the permuted set. As in our analysis with Gowinda, we extended genes by 1,000 base pairs in either direction before determining which genes contained a putative sweep in both the real and permuted SweepFinder2 scores.

### Analysis of the Vgsc region

*Vgsc* has long been studied in *Ae. aegypti* because of its relevance to insecticide resistance; therefore, we examined this gene and its surroundings. In addition to the genome-wide approach already detailed, we used bcftools mpileup and call, with the multiallelic variant calling model, to genotype variants present between 315 and 317 Mb of AaegL5_3, and on scaffold NIGP01000811. To visualize the relationship between scaffold NIGP01000811 and chromosome 3, we aligned the two using mummer 4.0.0rc1 (Marçais et al. 2018) and visualized the alignment using Ribbon (Nattestad et al. 2021).

We performed principal component analysis as previously described on all SNPs within the bounds of *Vgsc* (AAEL023266; AaegL5_3:315,926,360-316,405,639). In addition, we visualized the “loadings” of the first two principal components, plotted against each SNP’s chromosomal position. We also calculated nucleotide diversity (π) and Tajima’s *D* in the region spanning 310-320 Mb on chromosome 3, in 500 kb windows slid 50 kb.

In addition, we calculated linkage disequilibrium between individual variants, again using Rogers and Huff’s (2009) (Rogers and Huff 2009) estimate of *r*^2^, within *Vgsc* and 100 kb up- and downstream, after first masking low-quality genotypes as previously described. We calculated LD within all South American specimens, all North American specimens, and then all American specimens combined.

## DATA AND CODE AVAILABILITY

The nextflow pipelines used to align reads, call variants, and genotype variants can be found at https://github.com/rrlove/aedes_pipeline. Additional Python code used in data analysis can be found at <link>. Newly generated sequences from Colombia can be found at BioProject PRJNA864744.

## ACKNOWLEDGEMENTS

We thank Dr. Ben Matthews for sharing an *Aedes aegypti* genetic map. RRL and DRS were supported by the National Institutes of Health under award number R35GM138286. DRM was supported by National Institutes of Health under award number R01GM125715.

