## Supplemental figures for "Strong positive selection in *Aedes aegypti* and the rapid evolution of insecticide resistance"

### SUPPLEMENTAL FIGURE LEGENDS

**Figure S1.** Kinship coefficients between all 131 specimens.

**Figure S2.** Percent variance of the genome-wide PCA explained by each of the first ten principal components.

**Figure S3.** Distributions of nucleotide diversity π and Tajima's *D* by chromosome and country.

**Figure S4.** Nucleotide diversity π and Tajima's *D* along each chromosome, by country.

**Figure S5.** Distribution of linkage disequilibrium ($r^2$) by chromosome and country.

**Figure S6.** Composite likelihood ratio (CLR) as calculated by SweepFinder2, and nearby genes, for selected candidate sweeps. In each subfigure, the colored dots at the top of the main panel indicate individual test loci at 1 kb intervals that have sweep likelihood (CLR) values at or above the $99^{th}$ percentile for that country. Main panel line plots show CLR values averaged in 10 kb non-overlapping windows. Rectangles below the main panel indicate genes. Protein-coding genes with a predicted function are shown as white rectangles with black diagonal hatch marks in the upper half of the panel; all other genes are shown as transparent grey rectangles in the lower half of the panel. In subfigure B, the blue rectangles with the "x" hatch marks in the upper half of the panel mark glutathione transferase genes. In subfigure E, the blue rectangle with the “x” hatch marks in the lower half of the panel marks the voltage-sensitive sodium channel gene *Vgsc* (AAEL023266).

**Figure S7.** Functional enrichment of genes overlapping outlier sweep loci, across six cohorts. A 1 kb buffer was added up- and downstream of each gene before calculating enrichment. A black box at the intersection of a country column and GO term row indicates that term is enriched in genes overlapping outlier windows in that cohort.

**Figure S8.** Four populations show outlier windows overlapping a cluster of histone genes (red) on chromosome 3. Each panel labeled with a cohort shows individual outlier windows between 340.5 and 342.5 Mb on chromosome 3. Elevated set of red rectangles below the main plot show histone genes; all other genes are represented by grey rectangles.

**Figure S9.** Alignment depth and mapping quality on chromosome 3 from 310 to 320 Mb. Values were averaged across all specimens, and then averaged in 10 kb windows slid 1 kb.

**Figure S10.** Ribbon plot showing mummer alignment between NIGP01000811, and chromosome 3. Alignments shorter than 1 kb or with less than 90% identity between query and subject are excluded.

**Figure S11.** Principal component analysis of *Vgsc* (AAEL023266) and surrounding region using genotypes called with bcftools. PCA are shown with and without masking genotypes with a genotype quality less than 20.

**Figure S12.** Nucleotide diversity π from 310-320 Mb on AaegL5_3 for Brazilian and Colombian specimens in the top and bottom clusters of the *Vgsc* PCA. Values for entire Brazil and Colombia cohorts shown as grey solid and grey dashed lines, respectively. The vertical dashed lines mark the boundaries of *Vgsc* (315,926,360-316,405,639); the five Xs at the bottom indicate the locations of the five focal loci.

**Figure S13.** Nucleotide diversity π from 310-320 Mb on AaegL5_3 for specimens from Brazil, Colombia, and California, with the latter subdivided into three genetic clusters as in Lee et al. 2019. The vertical dashed lines mark the boundaries of *Vgsc* (315,926,360-316,405,639); the five Xs at the bottom of the Tajima's *D* plot indicate the locations of the five focal loci.

Figure S1

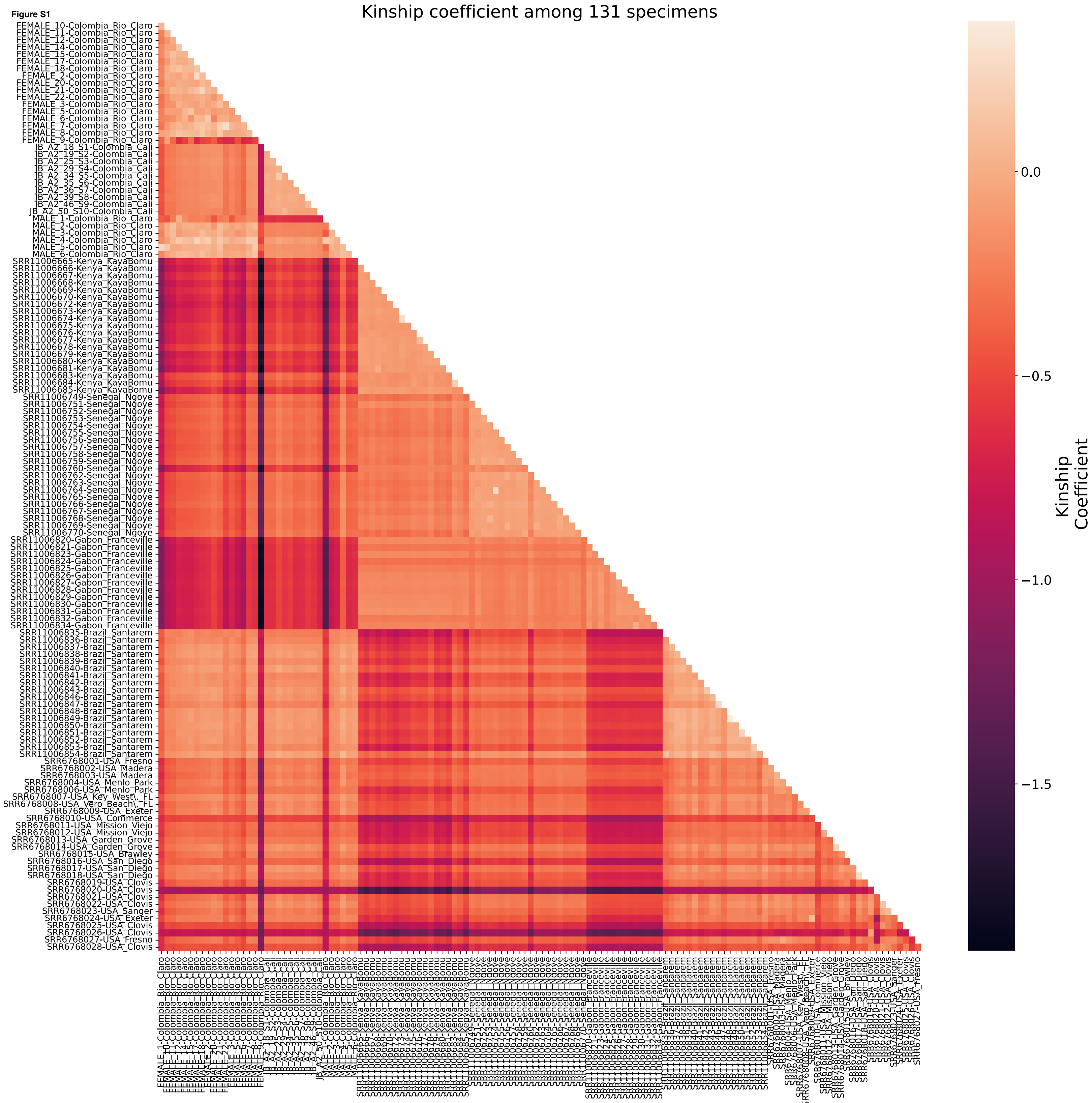

Kinship coefficient among 131 specimens
Kinship Coefficient
0.0
−0.5
−1.0
−1.5
FEMALE_10-Colombia_Rio_Claro
FEMALE_11-Colombia_Rio_Claro
FEMALE_12-Colombia_Rio_Claro
FEMALE_14-Colombia_Rio_Claro
FEMALE_15-Colombia_Rio_Claro
FEMALE_17-Colombia_Rio_Claro
FEMALE_18-Colombia_Rio_Claro
FEMALE_2-Colombia_Rio_Claro
FEMALE_20-Colombia_Rio_Claro
FEMALE_21-Colombia_Rio_Claro
FEMALE_22-Colombia_Rio_Claro
FEMALE_3-Colombia_Rio_Claro
FEMALE_5-Colombia_Rio_Claro
FEMALE_6-Colombia_Rio_Claro
FEMALE_7-Colombia_Rio_Claro
FEMALE_8-Colombia_Rio_Claro
FEMALE_9-Colombia_Rio_Claro
JB_A2_18_S1-Colombia_Cali
JB_A2_19_S2-Colombia_Cali
JB_A2_25_S3-Colombia_Cali
JB_A2_29_S4-Colombia_Cali
JB_A2_34_S5-Colombia_Cali
JB_A2_35_S6-Colombia_Cali
JB_A2_36_S7-Colombia_Cali
JB_A2_39_S8-Colombia_Cali
JB_A2_46_S9-Colombia_Cali
JB_A2_50_S10-Colombia_Cali
MALE_1-Colombia_Rio_Claro
MALE_2-Colombia_Rio_Claro
MALE_3-Colombia_Rio_Claro
MALE_4-Colombia_Rio_Claro
MALE_5-Colombia_Rio_Claro
MALE_6-Colombia_Rio_Claro
SRR11006665-Kenya_KayaBomu
SRR11006666-Kenya_KayaBomu
SRR11006667-Kenya_KayaBomu
SRR11006668-Kenya_KayaBomu
SRR11006669-Kenya_KayaBomu
SRR11006670-Kenya_KayaBomu
SRR11006672-Kenya_KayaBomu
SRR11006673-Kenya_KayaBomu
SRR11006674-Kenya_KayaBomu
SRR11006675-Kenya_KayaBomu
SRR11006676-Kenya_KayaBomu
SRR11006677-Kenya_KayaBomu
SRR11006678-Kenya_KayaBomu
SRR11006679-Kenya_KayaBomu
SRR11006680-Kenya_KayaBomu
SRR11006681-Kenya_KayaBomu
SRR11006683-Kenya_KayaBomu
SRR11006684-Kenya_KayaBomu
SRR11006685-Kenya_KayaBomu
SRR11006749-Senegal_Ngoye
SRR11006751-Senegal_Ngoye
SRR11006752-Senegal_Ngoye
SRR11006753-Senegal_Ngoye
SRR11006754-Senegal_Ngoye
SRR11006755-Senegal_Ngoye
SRR11006756-Senegal_Ngoye
SRR11006757-Senegal_Ngoye
SRR11006758-Senegal_Ngoye
SRR11006759-Senegal_Ngoye
SRR11006760-Senegal_Ngoye
SRR11006762-Senegal_Ngoye
SRR11006763-Senegal_Ngoye
SRR11006764-Senegal_Ngoye
SRR11006765-Senegal_Ngoye
SRR11006766-Senegal_Ngoye
SRR11006767-Senegal_Ngoye
SRR11006768-Senegal_Ngoye
SRR11006769-Senegal_Ngoye
SRR11006770-Senegal_Ngoye
SRR11006820-Gabon_Franceville
SRR11006821-Gabon_Franceville
SRR11006823-Gabon_Franceville
SRR11006824-Gabon_Franceville
SRR11006825-Gabon_Franceville
SRR11006826-Gabon_Franceville
SRR11006827-Gabon_Franceville
SRR11006828-Gabon_Franceville
SRR11006829-Gabon_Franceville
SRR11006830-Gabon_Franceville
SRR11006831-Gabon_Franceville
SRR11006832-Gabon_Franceville
SRR11006834-Gabon_Franceville
SRR11006835-Brazil_Santarem
SRR11006836-Brazil_Santarem
SRR11006837-Brazil_Santarem
SRR11006838-Brazil_Santarem
SRR11006839-Brazil_Santarem
SRR11006840-Brazil_Santarem
SRR11006841-Brazil_Santarem
SRR11006842-Brazil_Santarem
SRR11006843-Brazil_Santarem
SRR11006846-Brazil_Santarem
SRR11006847-Brazil_Santarem
SRR11006848-Brazil_Santarem
SRR11006849-Brazil_Santarem
SRR11006850-Brazil_Santarem
SRR11006851-Brazil_Santarem
SRR11006852-Brazil_Santarem
SRR11006853-Brazil_Santarem
SRR11006854-Brazil_Santarem
SRR6768001-USA_Fresno
SRR6768002-USA_Madera
SRR6768003-USA_Madera
SRR6768004-USA_Menlo_Park
SRR6768006-USA_Menlo_Park
SRR6768007-USA_Key_West\,_FL
SRR6768008-USA_Vero_Beach\,_FL
SRR6768009-USA_Exeter
SRR6768010-USA_Commerce
SRR6768011-USA_Mission_Viejo
SRR6768012-USA_Mission_Viejo
SRR6768013-USA_Garden_Grove
SRR6768014-USA_Garden_Grove
SRR6768015-USA_Brawley
SRR6768016-USA_San_Diego
SRR6768017-USA_San_Diego
SRR6768018-USA_San_Diego
SRR6768019-USA_Clovis
SRR6768020-USA_Clovis
SRR6768021-USA_Clovis
SRR6768022-USA_Clovis
SRR6768023-USA_Sanger
SRR6768024-USA_Exeter
SRR6768025-USA_Clovis
SRR6768026-USA_Clovis
SRR6768027-USA_Fresno
SRR6768028-USA_Clovis

**Figure S2**

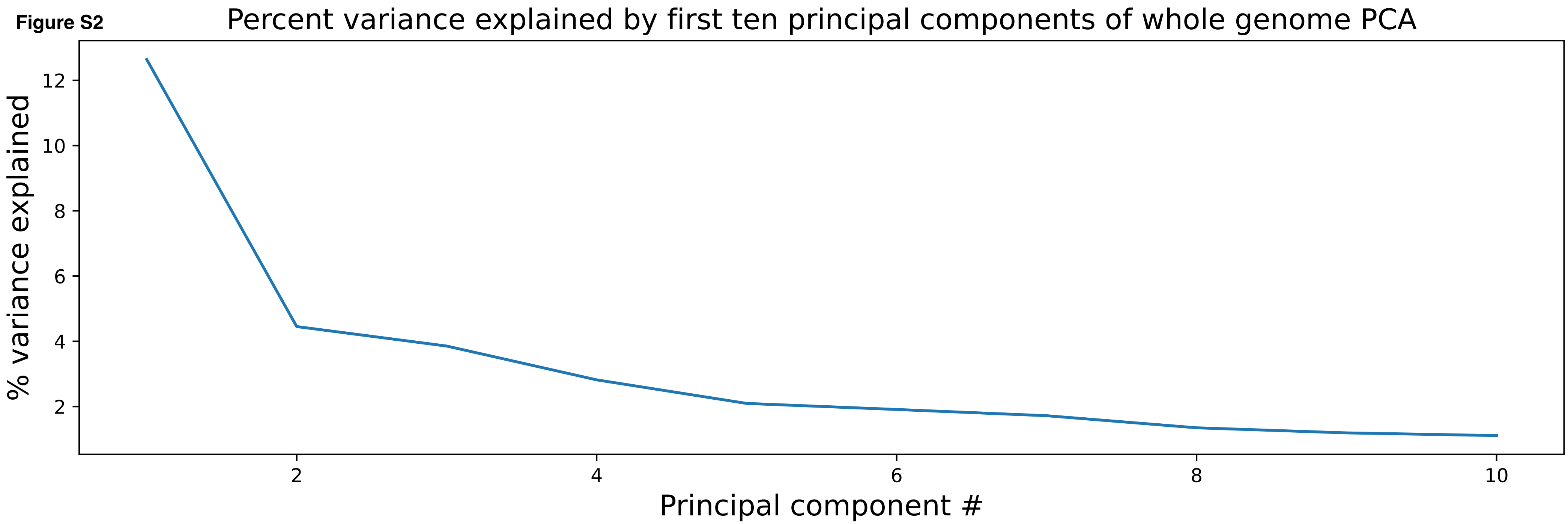
Percent variance explained by first ten principal components of whole genome PCA
% variance explained
2
4
6
8
10
12
Principal component #
2
4
6
8
10

Figure S3

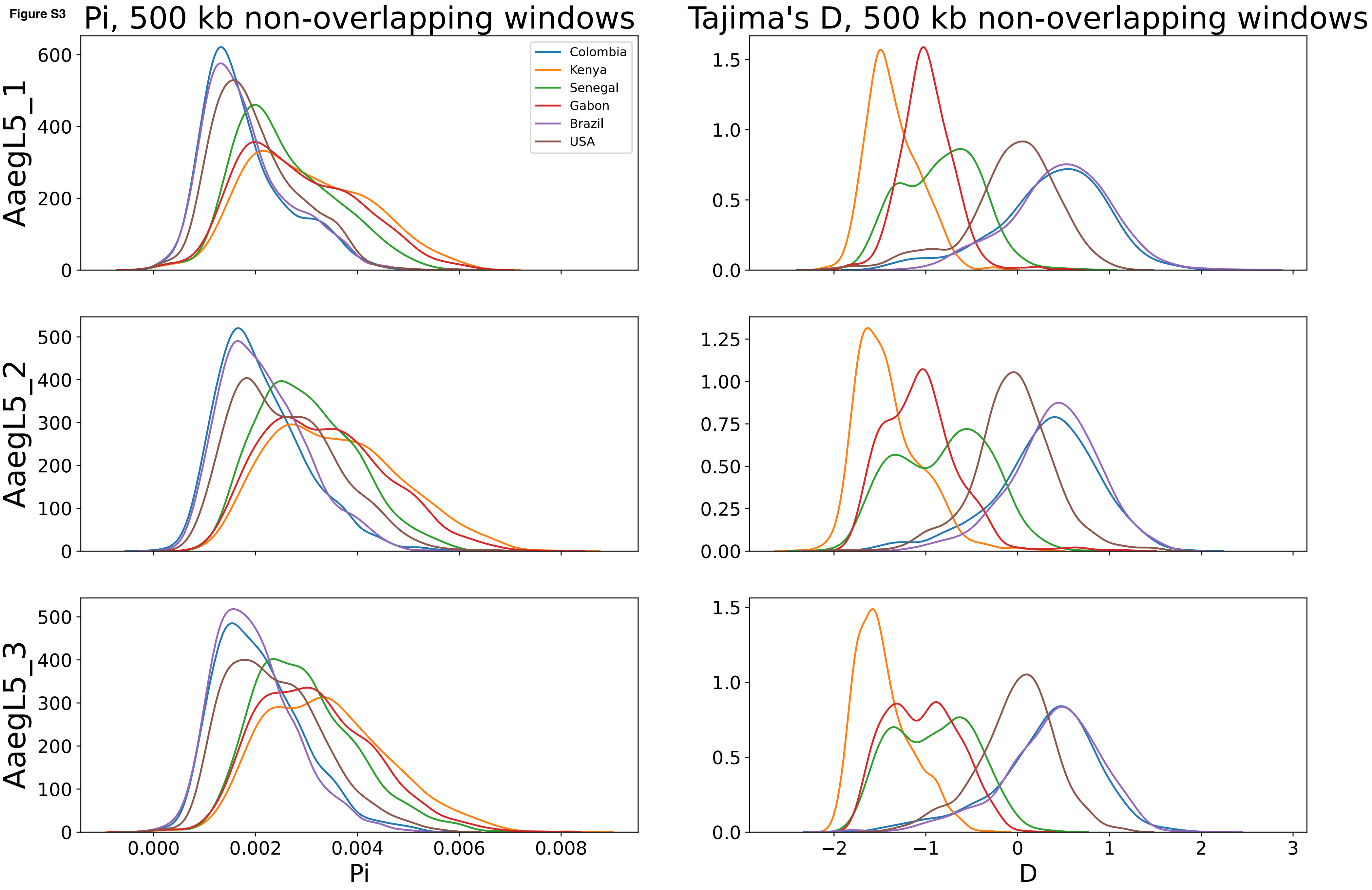

Pi, 500 kb non-overlapping windows
Tajima's D, 500 kb non-overlapping windows
Colombia
Kenya
Senegal
Gabon
Brazil
USA
AaegL5_1
AaegL5_2
AaegL5_3
Pi
D

Figure S4

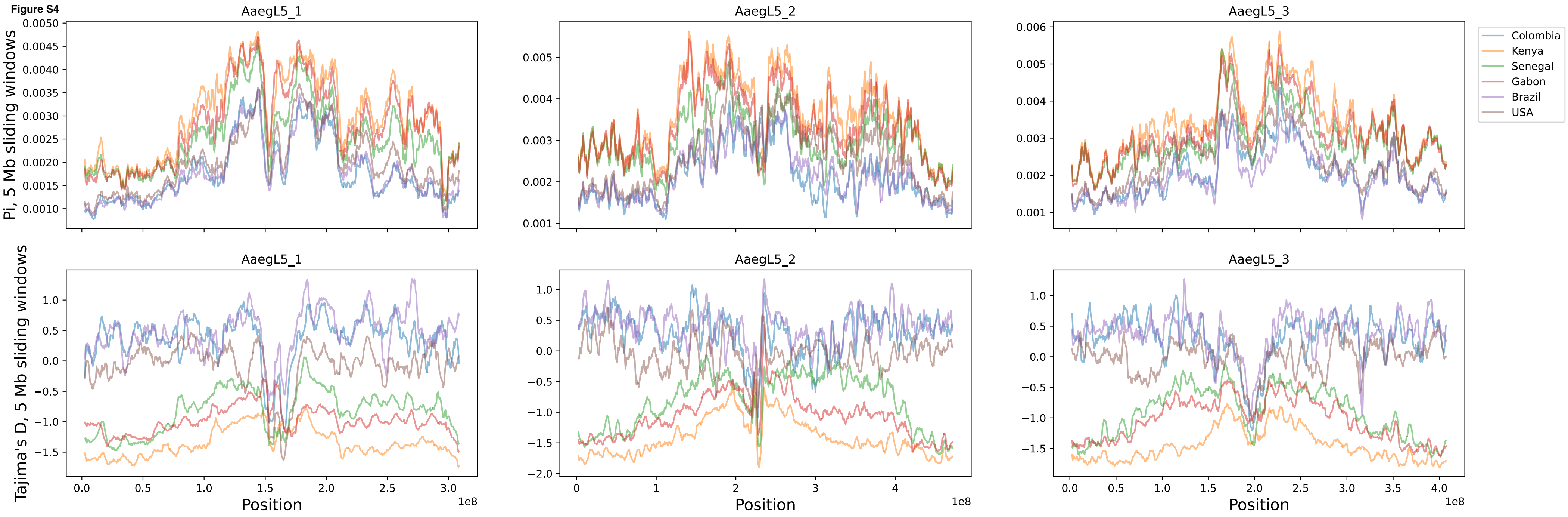
AaegL5_1
AaegL5_2
AaegL5_3
Pi, 5 Mb sliding windows
Tajima's D, 5 Mb sliding windows
Position
1e8
Colombia
Kenya
Senegal
Gabon
Brazil
USA

Figure S5

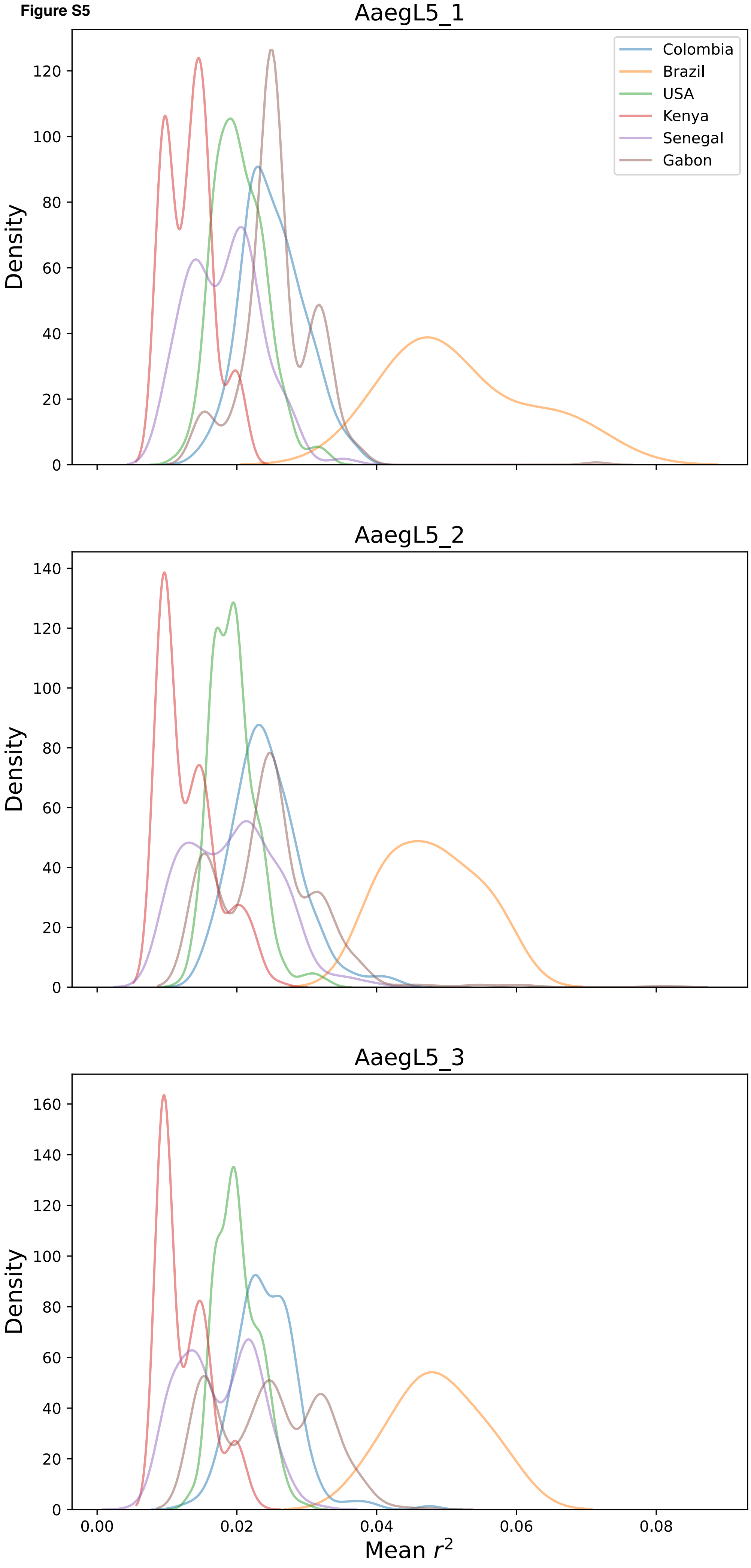

AaegL5_1
Colombia
Brazil
USA
Kenya
Senegal
Gabon
Density
0
20
40
60
80
100
120
AaegL5_2
Density
0
20
40
60
80
100
120
140
AaegL5_3
Density
0
20
40
60
80
100
120
140
160
0.00
0.02
0.04
0.06
0.08
Mean $r^2$

Figure S6

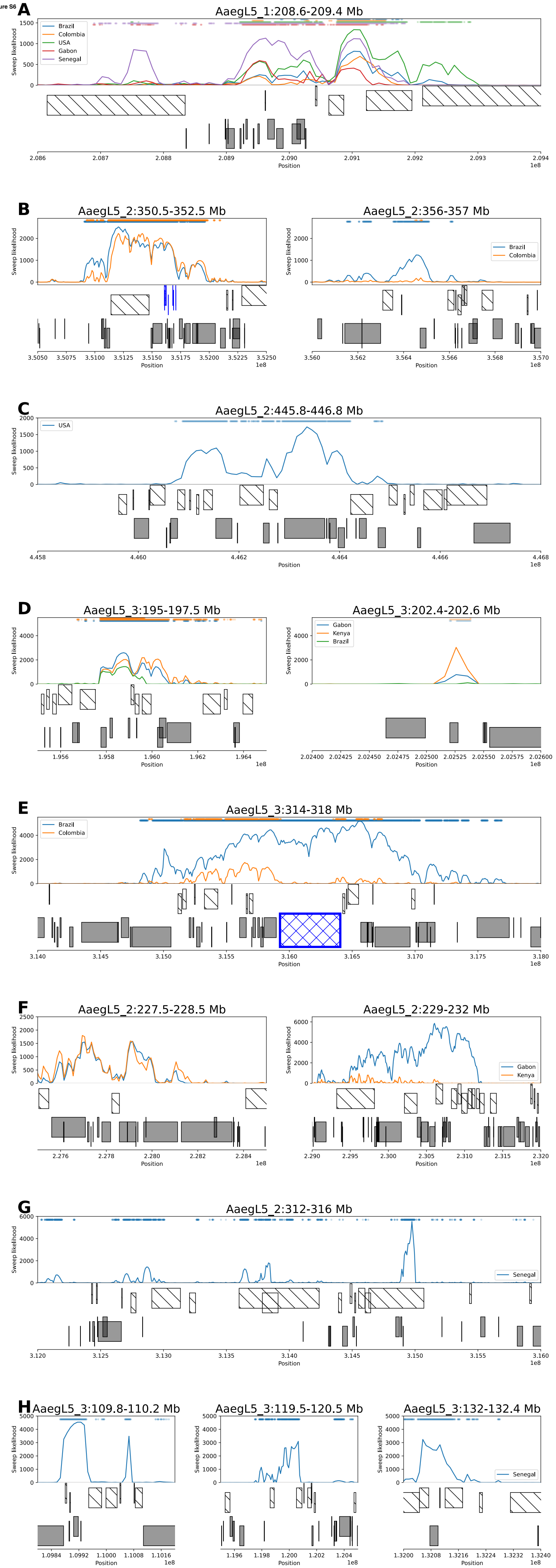
A
AaegL5_1:208.6-209.4 Mb
Brazil
Colombia
USA
Gabon
Senegal
Sweep likelihood
Position
1e8
B
AaegL5_2:350.5-352.5 Mb
AaegL5_2:356-357 Mb
Brazil
Colombia
C
AaegL5_2:445.8-446.8 Mb
USA
D
AaegL5_3:195-197.5 Mb
AaegL5_3:202.4-202.6 Mb
Gabon
Kenya
Brazil
E
AaegL5_3:314-318 Mb
Brazil
Colombia
F
AaegL5_2:227.5-228.5 Mb
AaegL5_2:229-232 Mb
Gabon
Kenya
G
AaegL5_2:312-316 Mb
Senegal
H
AaegL5_3:109.8-110.2 Mb
AaegL5_3:119.5-120.5 Mb
AaegL5_3:132-132.4 Mb
Senegal

**Figure S7**

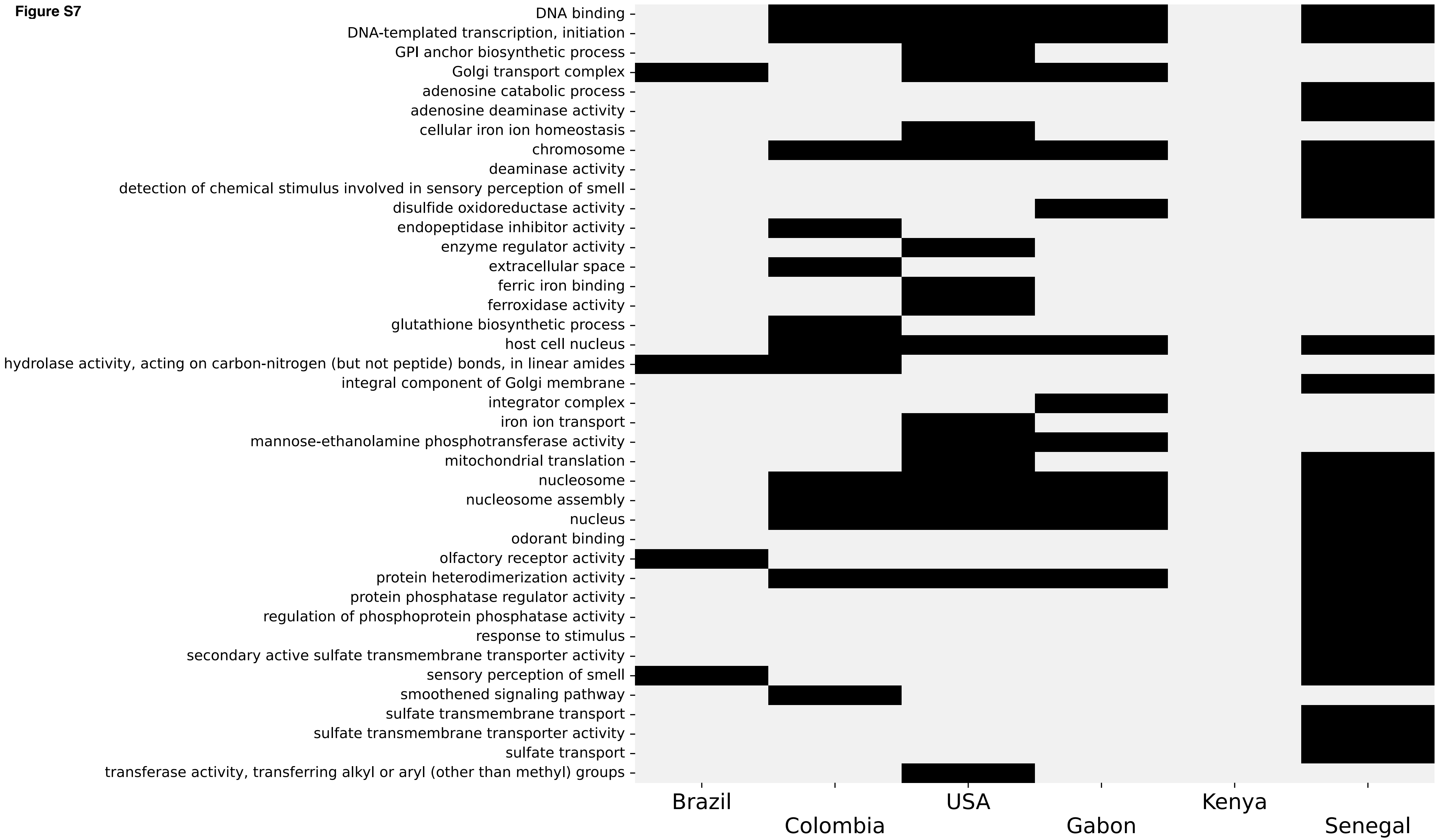
DNA binding
DNA-templated transcription, initiation
GPI anchor biosynthetic process
Golgi transport complex
adenosine catabolic process
adenosine deaminase activity
cellular iron ion homeostasis
chromosome
deaminase activity
detection of chemical stimulus involved in sensory perception of smell
disulfide oxidoreductase activity
endopeptidase inhibitor activity
enzyme regulator activity
extracellular space
ferric iron binding
ferroxidase activity
glutathione biosynthetic process
host cell nucleus
hydrolase activity, acting on carbon-nitrogen (but not peptide) bonds, in linear amides
integral component of Golgi membrane
integrator complex
iron ion transport
mannose-ethanolamine phosphotransferase activity
mitochondrial translation
nucleosome
nucleosome assembly
nucleus
odorant binding
olfactory receptor activity
protein heterodimerization activity
protein phosphatase regulator activity
regulation of phosphoprotein phosphatase activity
response to stimulus
secondary active sulfate transmembrane transporter activity
sensory perception of smell
smoothened signaling pathway
sulfate transmembrane transport
sulfate transmembrane transporter activity
sulfate transport
transferase activity, transferring alkyl or aryl (other than methyl) groups
Brazil
Colombia
USA
Gabon
Kenya
Senegal

Figure S8

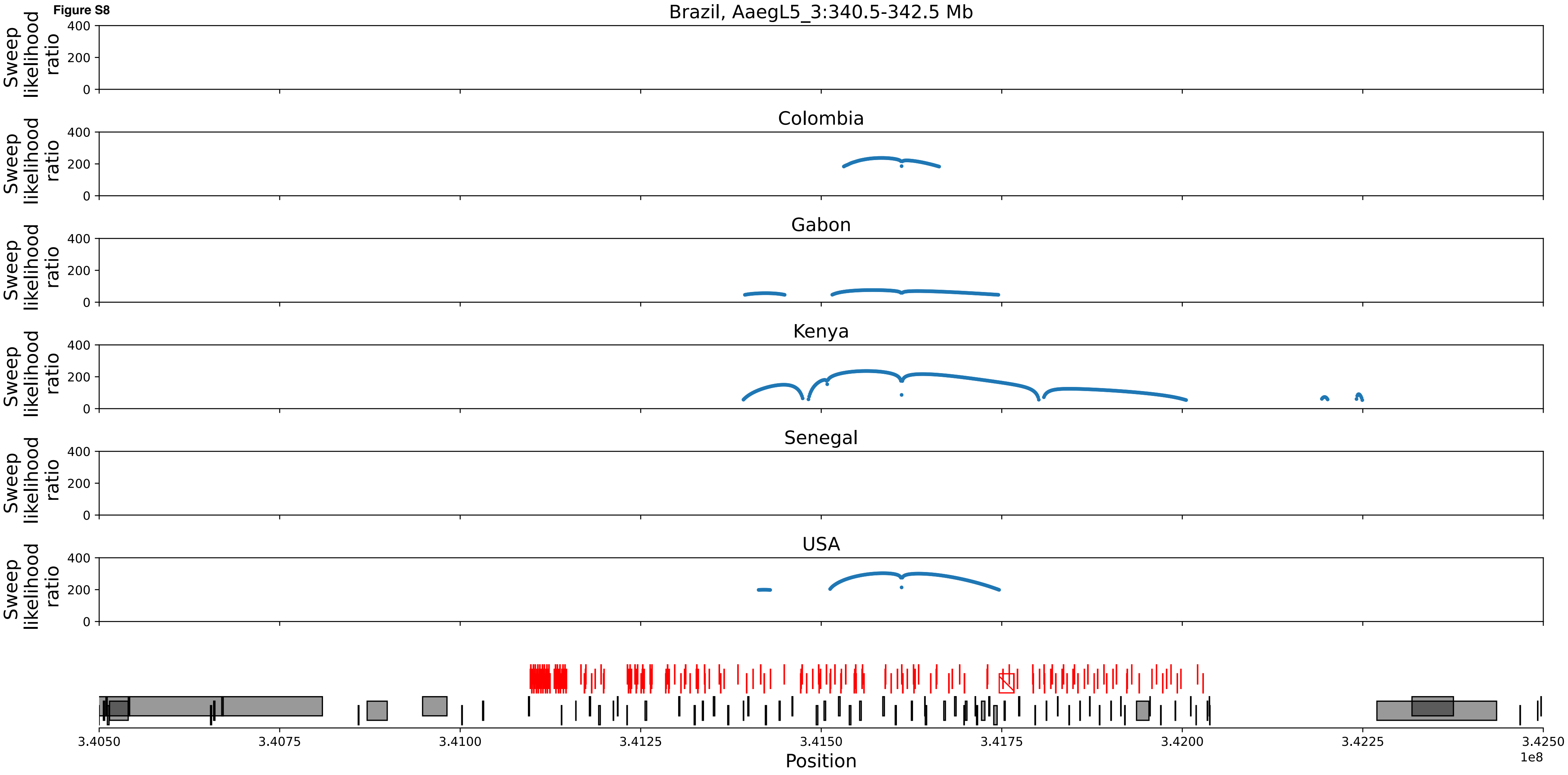
Brazil, AaegL5_3:340.5-342.5 Mb
Colombia
Gabon
Kenya
Senegal
USA
Sweep likelihood ratio
400
200
0
3.4050
3.4075
3.4100
3.4125
3.4150
3.4175
3.4200
3.4225
3.4250
1e8
Position

Figure S9

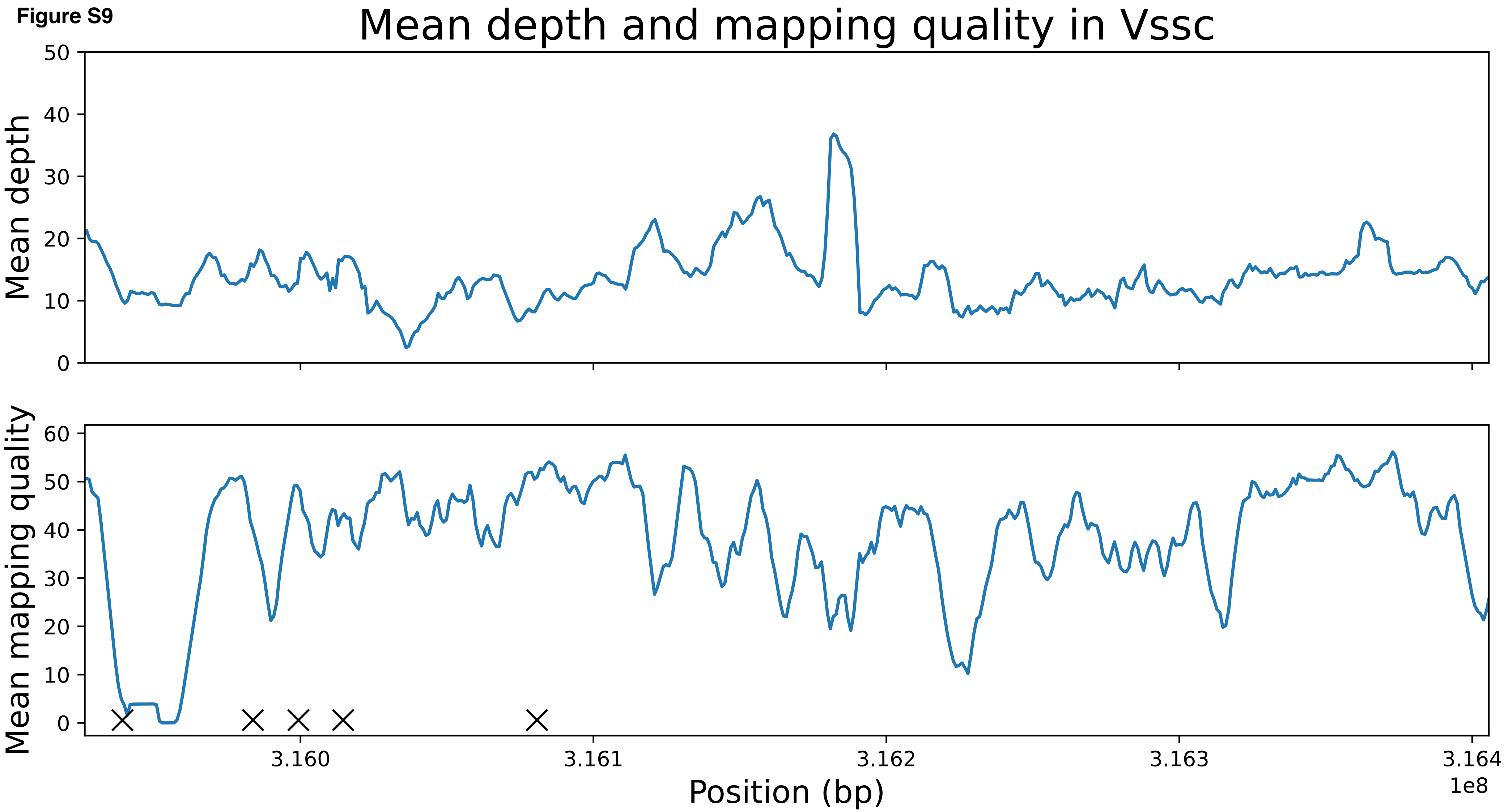
Mean depth and mapping quality in Vssc
Mean depth
0
10
20
30
40
50
Mean mapping quality
0
10
20
30
40
50
60
3.160
3.161
3.162
3.163
3.164
1e8
Position (bp)

**Figure S10**

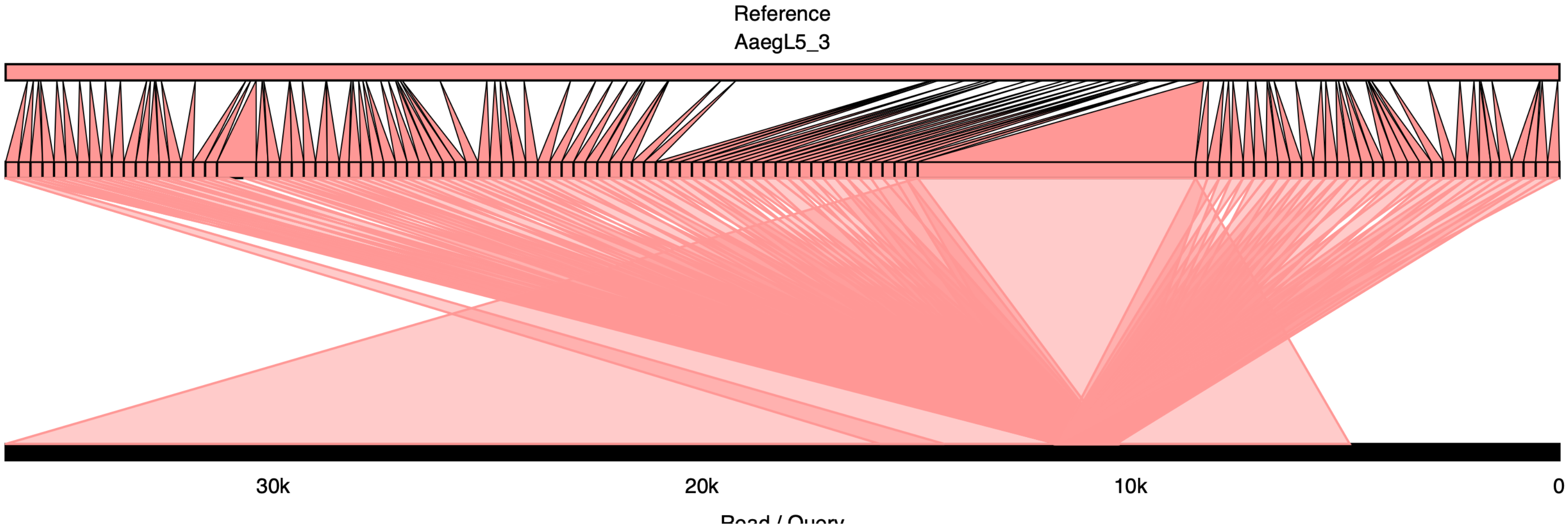

Figure S11

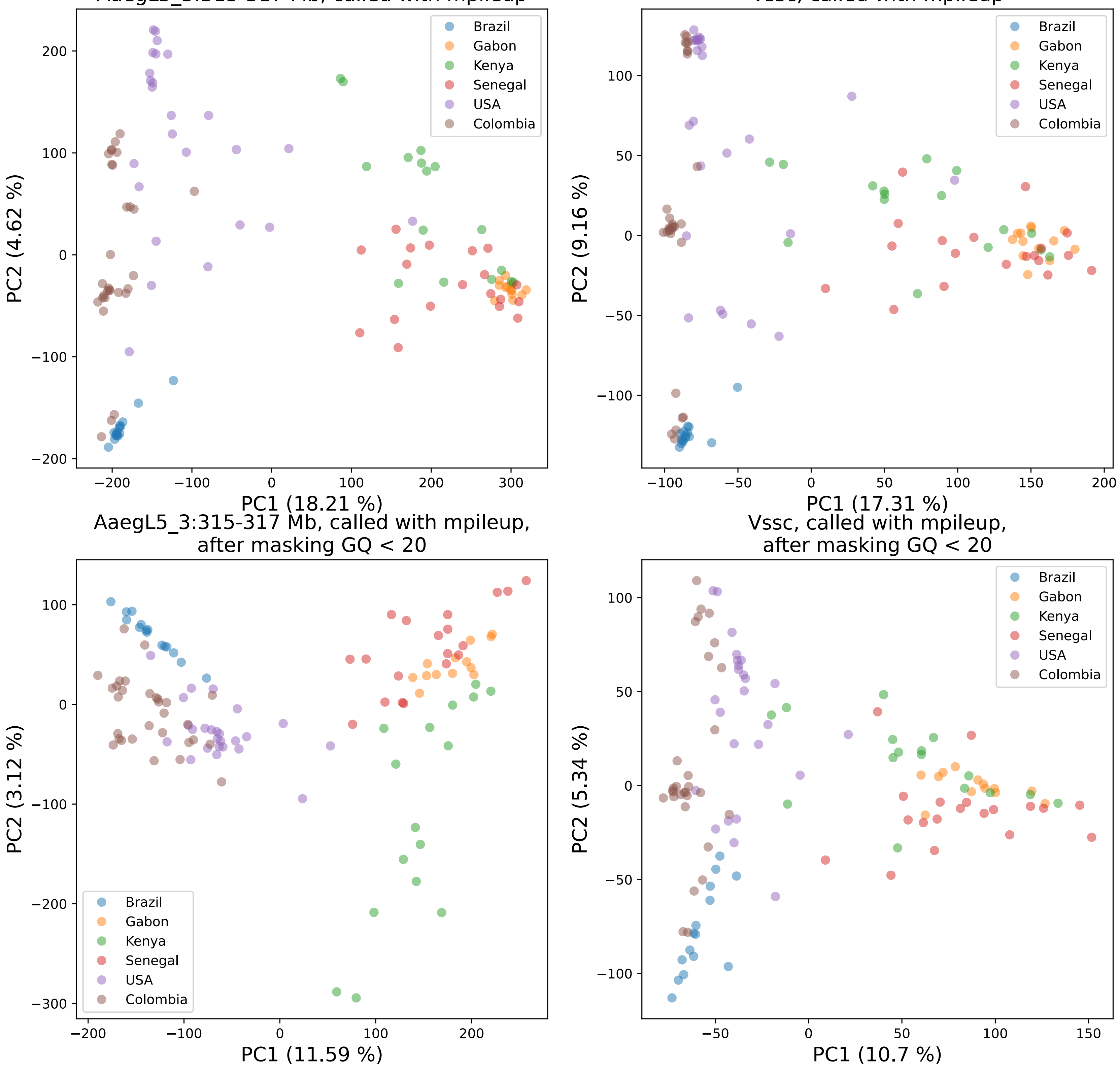
AaegL5_3:315-317 Mb, called with mpileup
PC1 (18.21 %)
PC2 (4.62 %)
Vssc, called with mpileup
PC1 (17.31 %)
PC2 (9.16 %)
AaegL5_3:315-317 Mb, called with mpileup, after masking GQ < 20
PC1 (11.59 %)
PC2 (3.12 %)
Vssc, called with mpileup, after masking GQ < 20
PC1 (10.7 %)
PC2 (5.34 %)
Brazil
Gabon
Kenya
Senegal
USA
Colombia

**Figure S12**

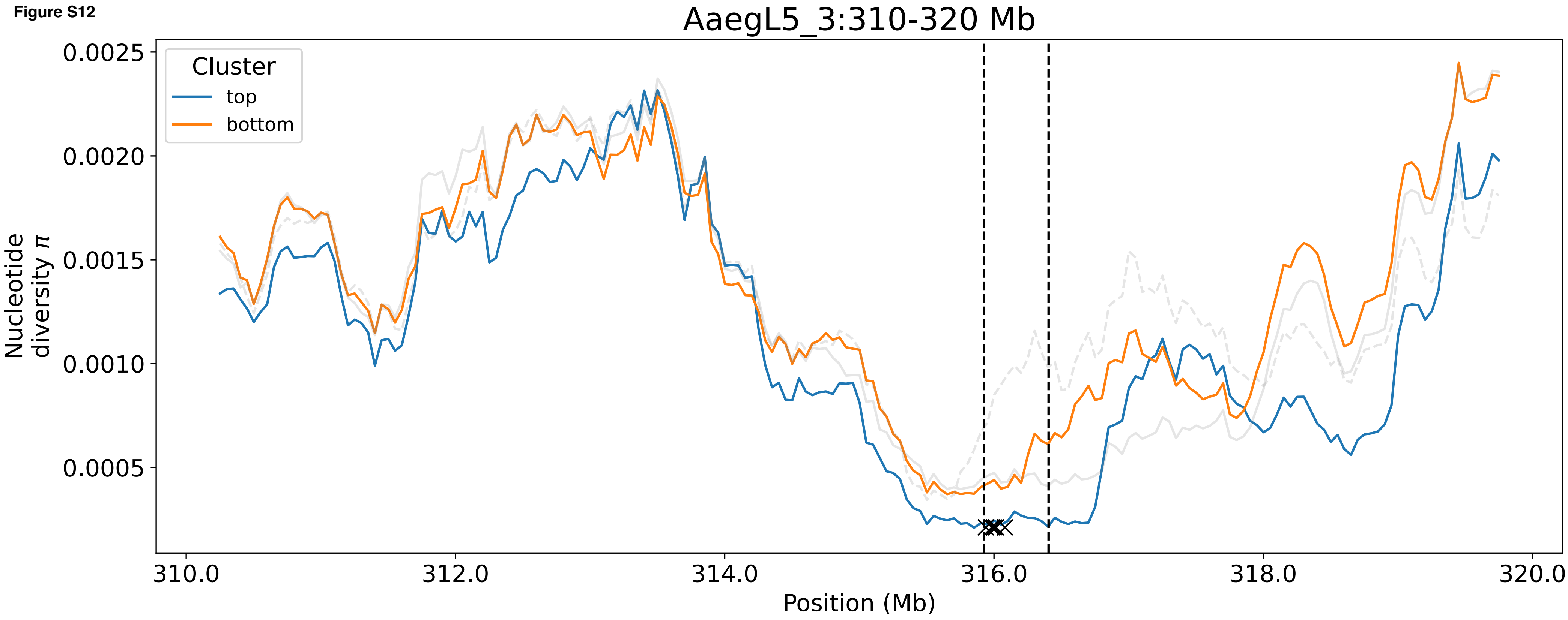

AaegL5_3:310-320 Mb
Cluster
top
bottom
Nucleotide diversity π
0.0025
0.0020
0.0015
0.0010
0.0005
310.0
312.0
314.0
316.0
318.0
320.0
Position (Mb)

**Figure S13**

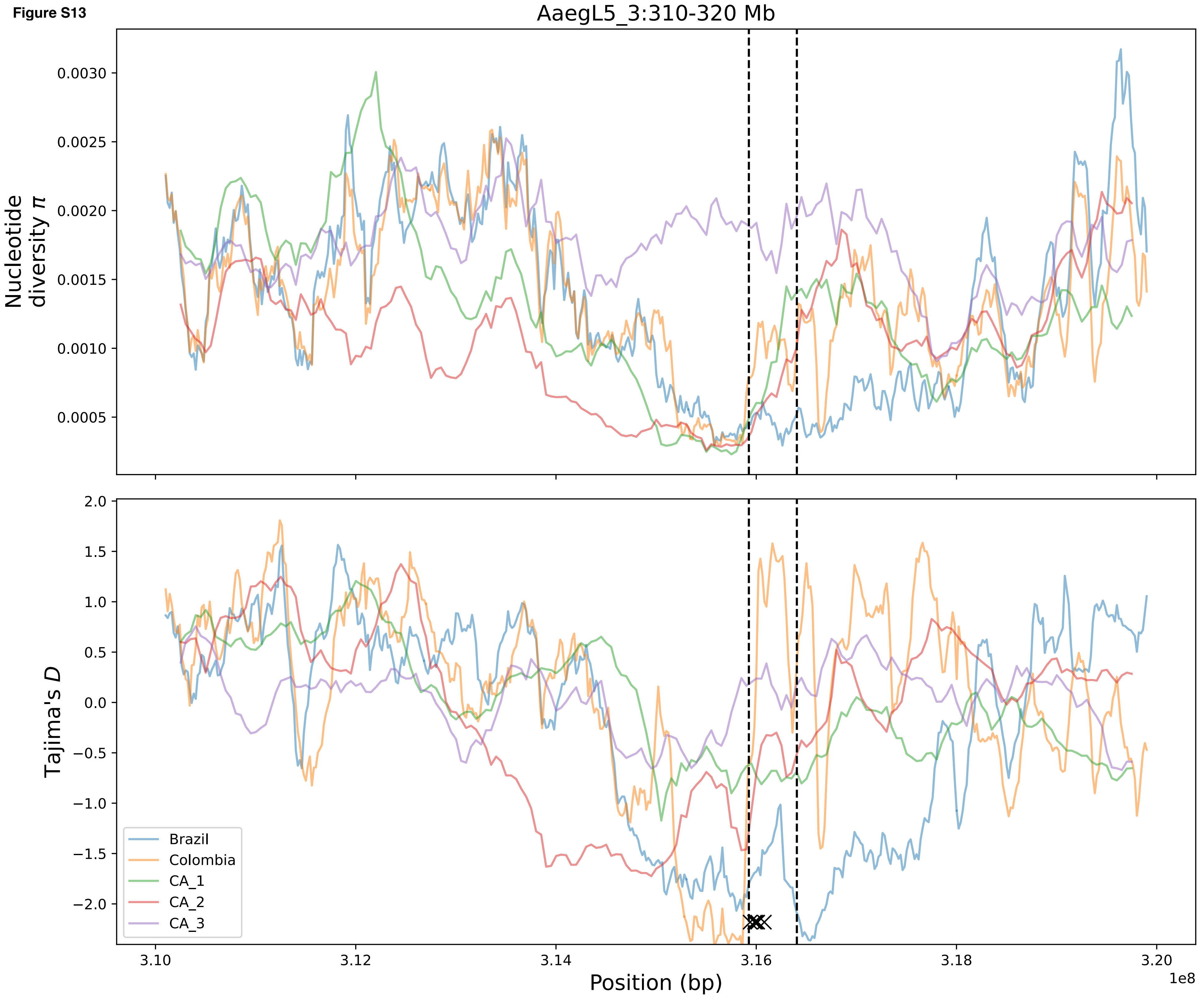
AaegL5_3:310-320 Mb
Nucleotide diversity π
0.0030
0.0025
0.0020
0.0015
0.0010
0.0005
Tajima's D
2.0
1.5
1.0
0.5
0.0
−0.5
−1.0
−1.5
−2.0
Brazil
Colombia
CA_1
CA_2
CA_3
3.10
3.12
3.14
3.16
3.18
3.20
1e8
Position (bp)
