## Supplemental tables 2-6, 8 for "Strong positive selection in *Aedes aegypti* and the rapid evolution of insecticide resistance"

Table S1. Specimen provenances (separate file).

Table S2. Alignment quality by country.

| | Mean (median) average read depth, per specimen | mean (median) % reads mapping, per specimen |
|---|---|---|
| Brazil | 13.06 (13.08) | 97.82 (97.89) |
| Colombia | 21.19 (20.90) | 96.69 (98.37) |
| USA | 10.16 (9.03) | 98.19 (98.19) |
| Gabon | 14.82 (14.94) | 94.80 (94.65) |
| Kenya | 15.64 (14.79) | 96.73 (96.82) |
| Senegal | 16.42 (15.80) | 96.94 (97.00) |

Table S3. Total and mean number of variants segregating in each cohort after removal of close kin. Columns normalized by cohort sample size have been rounded to the nearest SNP.

| | Total unfiltered SNP calls, before kin removal | Unfiltered SNP calls, per specimen, before kin removal | Total unfiltered SNP calls, after kin removal | Unfiltered SNP calls per specimen, after kin removal | Total filtered SNP calls | Filtered SNP calls, per specimen | Total SNP calls after kin removal | Per specimen |
|---|---|---|---|---|---|---|---|---|
| Brazil | 54,671,215 | 3,037,290 | 52,240,003 | 3,482,667 | 8,906,577 | 494,810 | 8,695,075 | 579,672 |
| Colombia | 71,992,561 | 2,117,428 | 70,728,776 | 2,357,626 | 10,619,884 | 312,350 | 10,505,852 | 350,195 |
| USA | 75,753,820 | 2,805,697 | 74,676,374 | 2,987,055 | 13,052,173 | 483,414 | 12,937,350 | 517,494 |
| Gabon | 99,034,712 | 7,618,055 | 99,034,712 | 7,618,055 | 19,220,350 | 1,478,488 | 19,220,350 | 1,478,488 |
| Kenya | 132,966,260 | 6,998,224 | 127,591,780 | 7,974,486 | 26,155,503 | 1,376,605 | 25,132,416 | 1,570,776 |
| Senegal | 106,893,209 | 5,344,660 | 105,765,723 | 5,566,617 | 18,818,551 | 940,928 | 18,657,154 | 981,955 |

Table S4. Nucleotide diversity and Tajima's *D* in six cohorts.

| Population | Mean (median) nucleotide diversity | Mean (median) Tajima's *D* |
|---|---|---|
| Brazil | 0.0020 (0.0019) | 0.406 (0.449) |
| Colombia | 0.0020 (0.0018) | 0.323 (0.390) |
| USA | 0.0024 (0.0022) | -0.040 (-0.009) |
| Gabon | 0.0032 (0.0030) | -1.031 (-1.036) |
| Kenya | 0.0034 (0.0033) | -1.409 (-1.470) |
| Senegal | 0.0029 (0.0028) | -0.850 (-0.820) |

Table S5. Mean (median) $F_{ST}$ calculated between the six countries in our combined dataset.

| | Brazil | Colombia | USA | Gabon | Kenya | Senegal |
|---|---|---|---|---|---|---|
| Brazil | | | | | | |
| Colombia | 0.073 (0.069) | | | | | |
| USA | 0.088 (0.090) | 0.091 (0.088) | | | | |
| Gabon | 0.238 (0.244) | 0.258 (0.265) | 0.190 (0.191) | | | |
| Kenya | 0.200 (0.197) | 0.220 (0.220) | 0.159 (0.163) | 0.062 (0.048) | | |
| Senegal | 0.136 (0.139) | 0.146 (0.143) | 0.104 (0.102) | 0.094 (0.083) | 0.107 (0.103) | |

Table S6. Mean (median) linkage disequilibrium by country across all three chromosomes, calculated in 500 kb nonoverlapping windows.

| Population | Mean (median) $r^2$ |
|---|---|
| Brazil | 0.049 (0.048) |
| Colombia | 0.025 (0.024) |
| USA | 0.020 (0.020) |
| Gabon | 0.025 (0.025) |
| Kenya | 0.013 (0.013) |
| Senegal | 0.018 (0.019) |

Table S7. Genes inside selected putative sweep regions (separate file).

Table S8. Linkage disequilibrium ($r^2$) between five known insecticide resistance loci, in the USA cohort.

| | F1534C (315939224) | V1016I (315983763) | I915K (315999297) | S723T (316014588) | V410L (316080722) |
|---|---|---|---|---|---|
| F1534C (315939224) | 1 | 0.278949 | 0.023957 | 0.162578 | 0.008951 |
| V1016I (315983763) | 0.278949 | 1 | 0.521999 | 0.537215 | 0.387558 |
| I915K (315999297) | 0.023957 | 0.521999 | 1 | 0.347025 | 0.404762 |
| S723T (316014588) | 0.162578 | 0.537215 | 0.347025 | 1 | 0.766312 |
| V410L (316080722) | 0.008951 | 0.387558 | 0.404762 | 0.766312 | 1 |
